# *De novo* phytosterol synthesis in animals

**DOI:** 10.1101/2022.04.22.489198

**Authors:** Dolma Michellod, Tanja Bien, Daniel Birgel, Marlene Jensen, Manuel Kleiner, Sarah Fearn, Caroline Zeidler, Harald R Gruber-Vodicka, Nicole Dubilier, Manuel Liebeke

## Abstract

Sterols are lipids that regulate multiple processes in eukaryotic cells, and are essential components of cellular membranes. Sterols are currently assumed to be kingdom specific, with phytosterol synthesis restricted to plants while animals are only able to synthesize cholesterol. Here, we challenge this assumption by demonstrating that the marine annelids *Olavius* and *Inanidrilus* synthesize the phytosterol sitosterol *de novo*. Using multi-omics, high-resolution metabolite imaging, heterologous gene expression and enzyme assays, we show that sitosterol is the most abundant (60%) sterol in these animals and characterize its biosynthetic pathway. We show that phytosterol synthesis partially overlaps with cholesterol synthesis and involves a non-canonical C-24 sterol methyltransferase (C_24_-SMT). C_24_-SMT is an essential enzyme for sitosterol synthesis in plants, but not known from animals with bilateral symmetry (bilaterians). Our comparative phylogenetic analyses of C_24_-SMT homologs revealed that these are widely distributed across annelids and other animal phyla, including sponges and rotifers. Our findings show that phytosterol synthesis and use is not restricted to the plant kingdom, and indicate that the evolution of sterols in animals is more complex than previously assumed.

## Introduction

Sterols are essential lipids present in all eukaryotes. They regulate the physical properties of biological membranes and are involved in the formation of specialized lipid-protein microdomains critical for signal transduction (Yeagle, 1985; Simons & Ikonen, 1997; Simons & Toomre, 2000). In addition, as precursors and cofactors of signaling molecules, they participate in many signaling and regulatory pathways (Chiang & Ferrell, 2020; Radhakrishnan et al., 2020; Sarkar & Chattopadhyay, 2022; Thummel & Chory, 2002). While sterols are ubiquitous in eukaryotes, their distribution is assumed to be kingdom specific: fungi synthesize ergosterol (C_28_); plants harbor a mixture of phytosterols (C_28_ to C_29_) dominated by sitosterol, stigmasterol, and campesterol (Lagarda et al., 2006) and animals use cholesterol (C_27_). These inter-kingdom differences reflect a complex evolutionary history of sterol synthesis. Previous phylogenetic analyses suggest that most enzymes for the biosynthesis of plant, fungal, and animal sterols were already present in the last eukaryotic common ancestor (LECA) (Desmond & Gribaldo, 2009; Summons et al., 2006), with the kingdom-specific distribution observed in extant species then evolving from LECA through multiple events of enzyme losses and specializations.

Kingdom-specific sterols differ from each other in only small structural details. For example, phytosterols differ from cholesterol by the presence of an extra methyl- or ethyl-group at position C_24._ This methylation is catalyzed by C_24_ sterol methyltransferase (C_24_-SMT), an enzyme widely distributed in plants, protists and fungi but absent in nearly all animals (Haubrich et al., 2015; Volkman, 2005). The only exception are some marine sponges, in which C_24_-SMT homologues are not related to phytosterol synthesis but assumed to participate in the synthesis of 24-isopropylcholesterol, a cholesterol derivative that serves as a sponge biomarker (Germer et al., 2017; Gold et al., 2016).

As with most other sterol enzymes, an ancestral C_24_-SMT was likely present in the LECA but lost early in the evolution of animals, probably after the divergence of sponges, explaining why this enzyme is absent from bilaterians and why animals lack the ability to produce phytosterols (Desmond & Gribaldo, 2009; Gold et al., 2016; Haubrich et al., 2015).

Here, we show that phytosterol biosynthesis and use is not restricted to plants. By screening multiple members of a globally distributed group of marine gutless annelids from the genera *Olavius* and *Inanidrilus*, we show that all eight species analyzed have sitosterol as their main sterol and express the genes needed for its synthesis. We characterized the sitosterol biosynthetic pathway used by these annelids and demonstrate the importance of a previously uncharacterized group of C_24_-SMT homologs encoded in these animals’ genomes. Using heterologous gene expression and enzymatic assays, we show that two animal C_24_-SMT homologs are functional and catalyzes both C-24 and C-28 methylations. Finally, we discovered that C_24_-SMT homologs are widely distributed across annelids and other animal phyla, including sponges and rotifers, suggesting that phytosterols may also be synthesized by other animals Our findings demonstrate that animals are capable of synthesizing and using phytosterol and that these molecules are not restricted to the plant kingdom, and suggest that the use of phytosterols as geological biomarkers for plants should be reconsidered

## Results and discussion

### Sitosterol is the main sterol in the marine gutless annelid *Olavius algarvensis*

*O. algarvensis* belongs to a group of gutless marine annelids that are found worldwide, mainly in coral-reef and seagrass sediments. These annelids lack a digestive system and are obligately associated with bacterial endosymbionts that provide them with nutrition (Giere, 1981, 1985; Kleiner et al., 2012; Woyke et al., 2006). As part of our ongoing efforts to characterize the *O. algarvensis* symbiosis, we analyzed the metabolome of single worm individuals using both gas chromatography-mass spectrometry (GC-MS) and high-performance liquid chromatography-mass spectrometry (HPLC-MS). These analyses revealed an unusual sterol composition, with sitosterol accounting for the majority of the sterols detected (60%), and the remainder consisting of cholesterol (**Figure 1A**). This was unexpected, as cholesterol usually dominates the sterol pool in bilaterians, often making up more than 90% of the total sterol content (Goad, 1981; Sissener et al., 2018). Sitosterol is a phytosterol and, among bilaterians, has only been reported as the most abundant sterol in a few phytoparasitic nematodes that are incapable of *de novo* sterol synthesis (Chitwood et al., 1985, 1987; Cole & Krusberg, 1967). In these plant parasites, it is unclear if the detected phytosterol is only present in the nematode gut content, or incorporated into their cells and tissue. The absence of a gut in *O. algarvensis* excludes sterol contamination from plant matter in the digestive tract. However, sitosterol could originate from the bacterial symbionts of *O. algarvensis*, which form a thick layer between the cuticle and the epidermis of the animal (**Figure 1B**).

**Figure 1.**
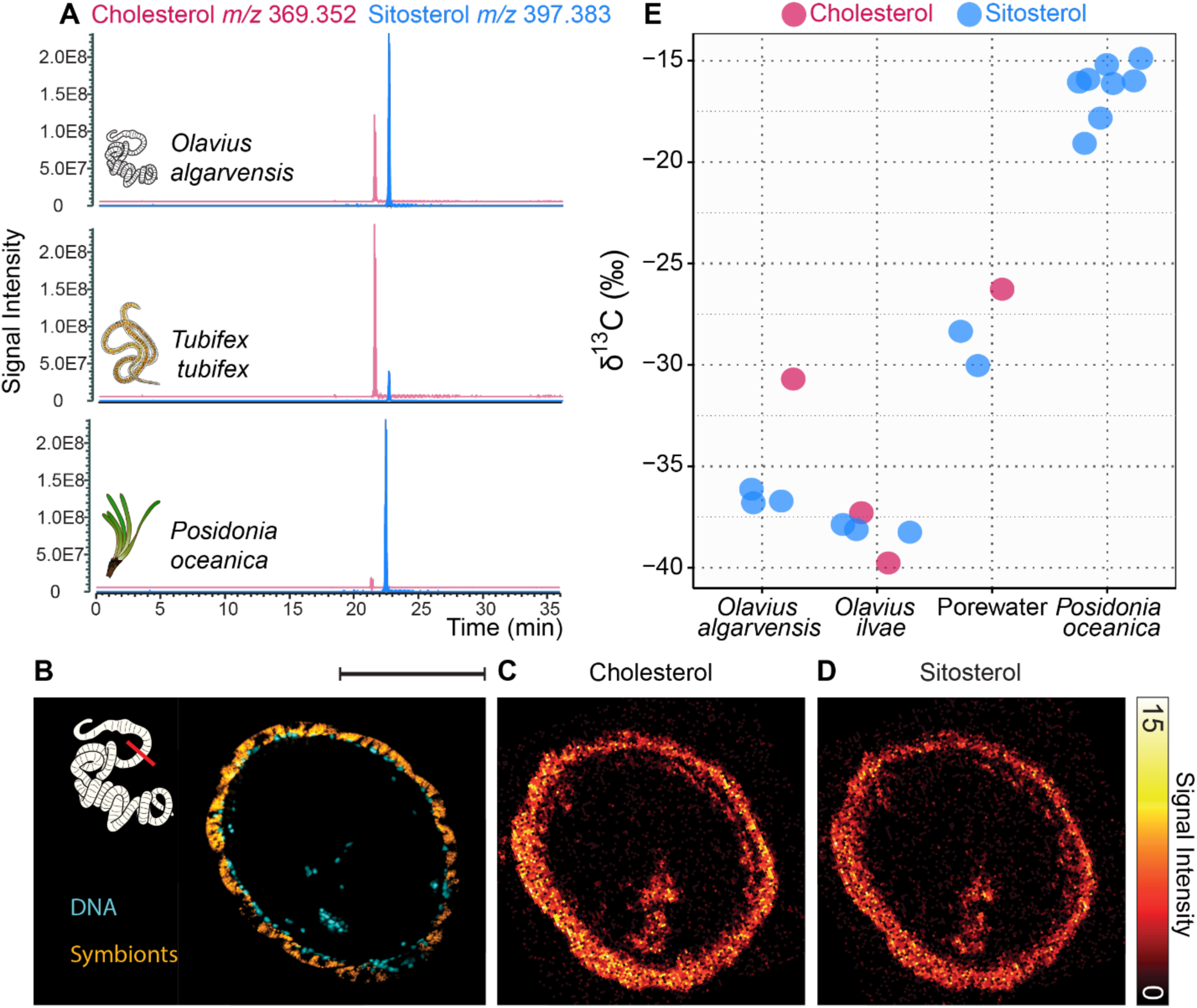
*Olavius algarvensis* has an unusual sterol profile dominated by sitosterol, a plant sterol. **A**, Extracted-ion chromatograms (XIC) of cholesterol ([M-H_2_O+H]^+^ C_27_H_45_ at *m/z* 369.352 (red)) and sitosterol ([M-H_2_O+H]^+^ C_29_H_49_ at *m/z* 397.383 (blue)). The XICs were generated from lipid extracts of (from top to bottom): the gutless marine annelid *O. algarvensis*, the freshwater annelid *Tubifex tubifex*, and the seagrass *Posidonia oceanica*. **B**, Chemosynthetic symbiotic bacteria are located just below the cuticle of *O. algarvensis*.16S rRNA fluorescence *in situ* hybridization (FISH) image of a cross section through a worm showing the symbionts in yellow (general eubacterial probe) and host nuclei in blue (DAPI). **C and D**, Distribution of sitosterol and cholesterol in *O. algarvensis* measured by TOF-SIMS **C**, Summed intensity of cholesterol ions (*m/z* 369.38, 385.34, 401.35) measured with TOF-SIMS. **D**, Summed intensity of sitosterol ions (*m/z*: 397.47, 383.37, 413.45) as measured by TOF-SIMS. **E**, The ^13^C isotopic composition of sterols in gutless annelids differed from that of the neighboring seagrass (*P. oceanica*) and sediment porewater. Scale bar in **B-D** 100 μm.

To investigate this possibility, we used two high spatial-resolution metabolite imaging techniques to localize the two major sterols in *O. algarvensis*. Time-of-Flight Secondary Ion Mass Spectrometry (TOF-SIMS) data revealed that, at a spatial resolution of 0.4 μm, both sitosterol and cholesterol were uniformly distributed throughout the animals’ tissues (**Figure 1C and 1D**). We found no evidence for a tissue-specific distribution of these two sterols, that is, there was no correlation between symbiont location and sitosterol distribution. These findings are supported by a second mass spectrometry imaging method, matrix-assisted laser desorption ionization mass spectrometry imaging (MALDI-2-MSI), of cross and longitudinal sections at a spatial resolution of 5 μm. The MALDI imaging data of longitudinal worm sections confirmed a uniform distribution of sitosterol and cholesterol throughout the animal (**Supplementary Figure 1 and 2**) and the identity of these sterols (**Supplementary Table 1)**. This homogeneous sterol distribution sterols suggests that the bacterial symbionts are not the source for phytosterol in *O. algarvensis*.

### *O. algarvensis* sterols have an isotopic composition that is distinct from their environment

Having found no evidence based on sterol distribution for a bacterial origin of sitosterol in *O. algarvensis*, we investigated if these animals acquire their sterols from the environment. Chemical analyses of porewater collected in the vicinity of seagrass meadows, the habitat of many gutless annelids (including *O. algarvensis*), showed that sterols were present in the environment in concentrations sufficient to sustain the growth of small sterol-auxotrophic invertebrates (**Supplementary information and Supplementary Figure 3**). Therefore, we further investigated the origin of sterols in *O. algarvensis* by analyzing the carbon isotopic signature (δ^13^C) of sterols in the worms, their environment (which includes the seagrass *Posidonia oceanica*) and the porewater of the sediments these worms live in. Carbon isotopic signatures are used to reveal carbon sources and their paths through the food web. As a rule, the bulk δ^13^C values of animals reflect their dietary sources (0.5‰ to 2‰ difference) (McCutchan et al., 2003; Tiunov, 2007). The δ^13^C values of sterols are depleted in ^13^C relative to bulk biomass by 5‰ to 8‰ (Canuel et al., 1997; Hayes, 2018). Results from gas chromatography isotope ratio mass spectrometry (GC-IRMS) with single metabolite resolution showed that sitosterol in the seagrass and porewater had δ^13^C values ranging from −30‰ to −15‰ (**Figure 1E and Supplementary information**). The sterols in *O. algarvensis*, as well as those in another co-occurring gutless annelid species, *Olavius ilvae*, had much lower δ^13^C values: −38‰ to −36‰ for sitosterol and −40‰ to −31‰ for cholesterol (**Figure 1E**). The difference in the isotopic signature of sterols in both *Olavius* species and their environment excludes that these worms acquired sterols from their environment, and instead indicates an endogenous origin. *O algarvensis*, as all other *Olavius* and *Inanidrilus* species, derives all of its nutrition from its chemosynthetic bacterial symbionts, and this is reflected in its bulk isotopic composition with δ^13^C values of −30.6‰ (Kleiner et al., 2015). The ^13^C-depleted signatures of both cholesterol and sitosterol by 1 to 10‰ compared to bulk biomass in *O. algarvensis* and *O. ilvae* led us to hypothesize that these animals synthesize both sterols *de novo*, using organic carbon derived from their chemosynthetic symbionts.

### *O. algarvensis* encodes and expresses enzymes involved in sitosterol synthesis that overlap with those of cholesterol synthesis

Having ruled out an external, environmental source of sitosterol in *O. algarvensis*, we next investigated if the animals themselves can synthesize this phytosterol. To identify and characterize the biosynthetic pathways involved in sterol production, we sequenced and assembled the genome of *O. algarvensis* and analyzed metatranscriptomic and metaproteomic data to search for enzymes involved in *de novo* sterol synthesis, screening both *O. algarvensis* and its symbionts. These analyses revealed that the symbionts, as most bacteria, do not encode enzymes involved in sterol synthesis. The host, on the other hand, possessed the full enzymatic toolbox required for cholesterol synthesis, with homologs of the 11 enzymes present in the genome of *O. algarvensis* (**Supplementary Figure 4 and Supplementary Table 2**). The cholesterol biosynthesis pathway, starting with squalene, is a series of ten connected enzymatic reactions encoded by 11 genes (**Supplementary Figure 4 and Supplementary Table 3**). Homologs of all enzymes were transcribed (11 out of 11 enzymes) and many of the proteins were detected in the proteome of *O. algarvensis* (5 out of 11 proteins) (**Supplementary Figure 4 and Supplementary Tables 4 and 5**), indicating active expression of the genes involved in cholesterol synthesis. Phylogenetic analysis allowed us to assign each homolog to an ortholog group and thus to a potential function (**Supplementary Figures 5 to 14**). Collectively, these data show that *O. algarvensis* has all the enzymes required for *de novo* cholesterol synthesis, which in combination with the isotopic signature of their cholesterol, suggests that these worms are able to synthesize cholesterol.

More importantly, our analyses also identified a homolog of C_24_-SMT, an enzyme essential to sitosterol synthesis, in the genome of *O. algarvensis* (**Figure 2**). As described above, bilaterians are thought to lack C_24_-SMT. C_24_-SMT catalyzes the transfer of a methyl group from S-adenosyl-L-methionine (SAM) to the sterol side chain and is essential for the biosynthesis of plant sterols. The gene structure of the putative C_24_-SMT confirmed its eukaryotic origin and excluded bacterial contamination (**Figure 2**). The putative C_24_-SMT gene is a 1071 bp open reading frame (ORF) encoding a 356 amino-acid polypeptide (**Supplementary Table 3**), and contains all the conserved residues characteristic of C_24_-SMT as well as the four conserved signature motifs responsible for substrate binding (**Supplementary Figure 15**) (Bouvier-Navé et al., 1998; Jayasimha & Nes, 2008; Nes et al., 2004, 2008; Nes & Heupel, 1986; Schaller, 2004; Veeramachaneni, P. P., 2005). We identified the C_24_-SMT gene in *O. algarvensis* transcriptomes and proteomes, confirming that these animals express this enzyme **(Figure 2, Supplementary Table 4 and 5**). Our findings suggest that the *O. algarvensis* C_24_-SMT gene encodes a functional enzyme that may be involved in sitosterol metabolism in these annelids, and represents the first discovery of a C_24_-SMT enzyme in bilaterians.

**Figure 2.**
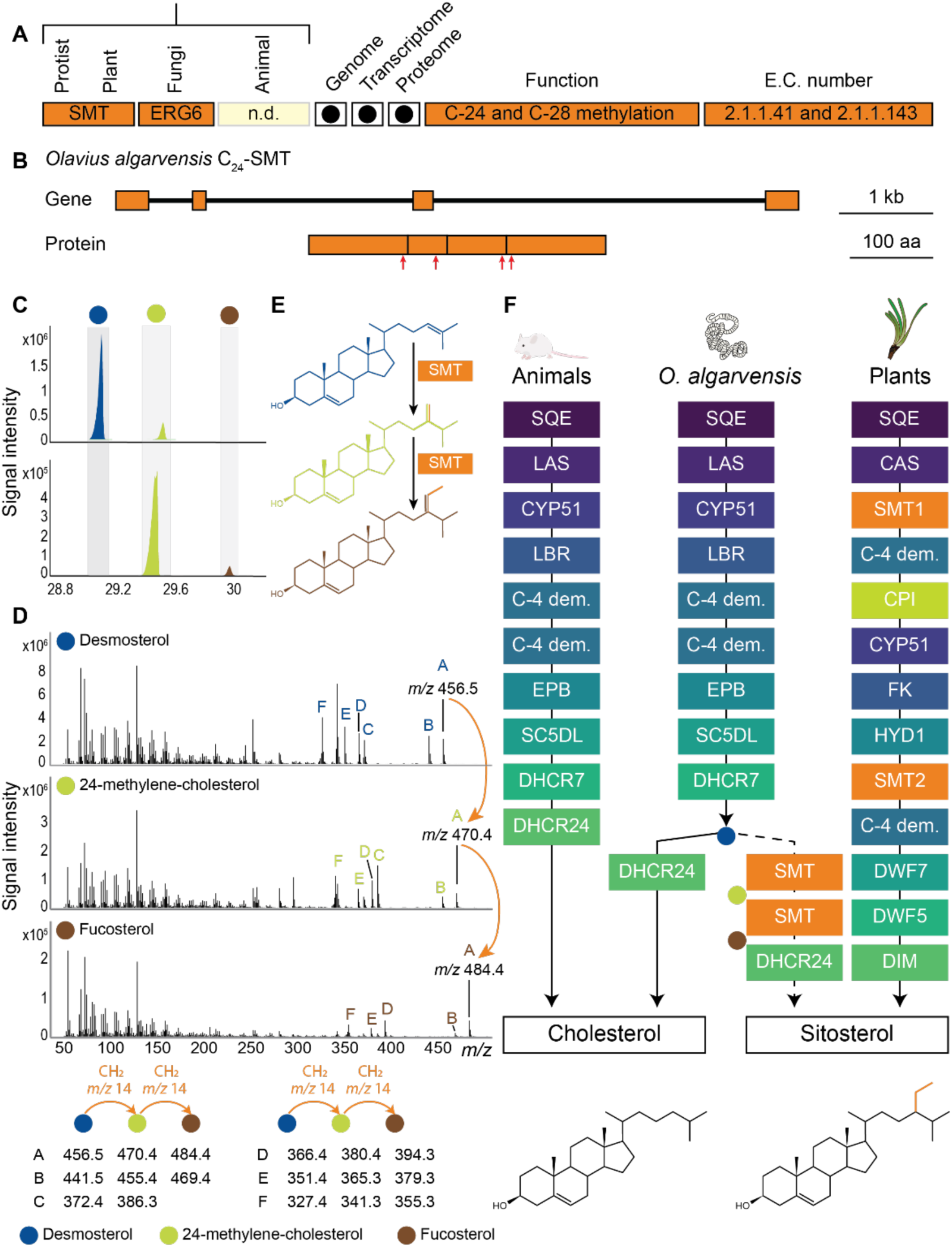
*Olavius algarvensis* encodes and expresses a C_24_-SMT which catalyzes two consecutive methylations, using desmosterol as the first and 24-methylene-cholesterol as the second substrate. **A**, *O. algarvensis* encodes and expresses a homolog of C_24_-SMT, an enzyme not previously found in bilaterian animals and essential to sitosterol synthesis. Dots indicate detection in the genome, transcriptome and proteome of *O. algarvensis*. **B**, The *O. algarvensis* C_24_-SMT gene consists of 4 exons forming a 1071 bp open reading frame encoding a 356 amino-acid polypeptide. The four conserved regions of the enzyme are highlighted by red arrows. **C**, Chromatograms of enzymatic assays with desmosterol (top) and 24-methylene-cholesterol (bottom) as substrates. *O. algarvensis* C_24_-SMT, after overexpression in *E. coli*, added a methyl group to the side chains of desmosterol and 24-methylene-cholesterol. In the first methylation desmosterol, an intermediate of cholesterol synthesis, was methylated to produce 24-methylene-cholesterol (C_28_ sterol). In the second methylation, 24-methylene-cholesterol was methylated to produce fucosterol (C_29_ sterol). **D**, Mass spectra of the different substrates and methylated products from the enzymatic assays. Sterol intermediates differ by the number of methyl groups (CH_2_ at *m/z* 14) attached to their side chain. The side chain of desmosterol is not methylated, 24-methylene-cholesterol has a methyl group at C-24, and fucosterol has two methyl groups at C-24 and C-28. The substrates and methylated products were identified by MS, retention time and comparison with standards. The fragmentation pattern suggests that the methyl groups were added to the side chain of the sterols. **E**, Structural representation of the two methylation steps in *O. algarvensis*. **F**, Comparison of the enzyme used in the proposed sterol synthesis pathways in *Olavius* to the canonical cholesterol and sitosterol synthesis pathways (similar enzymatic reactions are colored similarly). The first six steps are common to both cholesterol and sitosterol synthesis pathways. This trunk pathway branches off after the synthesis of desmosterol. For sitosterol synthesis, desmosterol is first methylated by C_24_-SMT to 24-methylene-cholesterol, which is then methylated in a second, consecutive step by C_24_-SMT to fucosterol. Fucosterol is reduced to sitosterol by a sterol C24-reductase (DHCR24, DIM). Squalene monooxygenase (SQE), oxydosqualene cyclase (LAS, CAS), sterol 14 demethylase (CYP51), sterol 14-reductase (LBR, FK), C-4 demethylation (C-4 dem.), Sterol Δ7-Δ8 isomerase (EBP, HYD1), sterol 5-desaturase (SC5DL, DWF7), sterol Δ7 reductase (DHCR7, DWF5), and C-24 sterol methyltransferase (C_24_-SMT, SMT1, SMT2).

### The *O. algarvensis* C_24_-SMT homolog is bifunctional and consecutively transfers methyl groups to sterol intermediates

Two methylation reactions are required for the final steps of sitosterol synthesis, one that adds a methyl group at C-24 and one at C-28. These reactions can be catalyzed by the same or different enzymes. For example, in most plants, C-24 and C-28 methylations are mediated by different enzyme isoforms, SMT1 and SMT2 (Bouvier-Navé et al., 1998; Hartmann, 2004; Neelakandan et al., 2009). Alternatively, in basal plants such as green algae, a single C_24_-SMT catalyzes both methylation steps, (Desmond & Gribaldo, 2009; Haubrich et al., 2015). Sterol profiles and genome analyses of sponges, where C_24_-SMTs participate in the synthesis of the cholesterol derivative 24-isopropylcholesterol, suggest that these enzymes are similarly bifunctional, although their biochemical activity remains to be characterized (Gold et al., 2016). Since *O. algarvensis* encodes a single C_24_-SMT homolog, we hypothesized that this enzyme mediates both C-24 and C-28 methylation.

To test this hypothesis, we overexpressed *O. algarvensis* C_24_-SMT in *Escherichia coli* and examined its enzymatic activity, substrate preferences, and products by assaying crude protein extracts with a methyl-donor (S-adenosylmethionine, SAM) and different sterol substrates. The C_24_-SMT from *O. algarvensis* expressed in *E. coli* was not able to methylate classical plant sterol substrates (**Supplementary Table 14)**. However, the enzyme was able to methylate zymosterol and desmosterol, two intermediates of the cholesterol biosynthetic pathway. When incubated with either of these sterol substrates and SAM, the *O. algarvensis* C_24_-SMT produced a methylated C_28_ product (**Figure 2 and Supplementary Figure 16-19**). Zymosterol was methylated to fecosterol and desmosterol to 24-methylene-cholesterol. The shift in retention times and changes in mass spectra of the products indicated that a methyl group was added to their sterol side chain, likely at the C_24_-position (**Supplementary Figure 18-19**). These results suggest that the cholesterol and sitosterol synthesis pathways overlap in *O. algarvensis*, as the two C_24_-SMT substrates, zymosterol and desmosterol, are intermediates produced in the second half of the animal cholesterol synthesis pathway (**Figure 2)**.

After confirming the first methylation step at C-24, we next searched for potential substrates for the second methylation step at C-28. This second methylation is essential as sitosterol is a C_29_ compound, characterized by the presence of two methyl groups on its side chain. To test our hypothesis that both of these methylations are catalyzed by the *O. algarvensis* C_24_-SMT, we selected the product of the first methylation, 24-methylene-cholestrol, as well as campesterol, as potential substrates for the second methylation (**Supplementary Figure 20**). 24-methylene-cholesterol was the only substrate to which the *O. algarvensis* C_24_-SMT added a methyl group, producing the C_29_ compound fucosterol (**Figure 2 and Supplementary Figure 20 and 21**). 24-methylene-cholesterol is the product of the methylation of desmosterol, providing evidence to support our hypothesis that in *O. algarvensis*, the C-24 and C-28 methylations are catalyzed by the same enzyme and occur consecutively. That is, the *O. algarvensis* C_24_- SMT first methylates desmosterol at C-24 to produce the C_28_ sterol 24-methylene-cholesterol, and then adds a second methyl group to 24-methylene-cholesterol at C-28, to produce the C_29_ sterol fucosterol. Fucosterol differs from sitosterol by the presence of a double bond at position C-24(28). This double bond is most likely removed by the delta(24)-sterol reductase (DHCR24), which is expressed based on its presence in *O. algarvensis* transcriptomes (**Supplementary Figure 4 and 14**). These results provide the first evidence for an animal C_24_-SMT able to catalyze the two methylation steps needed to synthesize sitosterol from a cholesterol intermediate, revealing a previously unknown pathway for phytosterol synthesis in animals (**Figure 2**).

### C_24_-SMT homologs are widespread in annelids

Having demonstrated the activity of an animal C_24_-SMT homolog that enables *O. algarvensis* to synthesize sitosterol *de novo*, we asked if other gutless annelids also encode functional C_24_-SMTs. To answer this question, we analyzed the sterol contents of six additional gutless annelid species, collected at locations in the Mediterranean Sea (Elba, Mallorca, Monaco) and the Caribbean Sea (Bahamas, Belize). All six species had similar lipid profiles as *O. algarvensis*, with sitosterol as their major sterol (**Supplementary Table 6 and Supplementary information**).

In addition to lipid profiling, we screened the transcriptomes of nine *Olavius* and *Inanidrilus* species and found all nine species expressed a C_24_-SMT homolog **(Supplementary Table 7)**, including *O. ilvae*, which had similarly negative sitosterol δ^13^C values as *O. algarvensis* (**Figure 1E**) and also encodes a C_24_-SMT in its genome (**Supplementary Table 3**). We confirmed the C-24 methylation ability of a second C_24_- SMT homolog, from *O. clavatus*, by heterologously expressing this gene in *E. coli*. Our biochemical assays demonstrated that the C_24_-SMT from *O. clavatus* is also a bifunctional sterol methyltransferase, capable of methylating zymosterol, desmosterol and 24-methylene-cholesterol (**Supplementary Figure 16, 17 and 21**).

We next asked if C_24_-SMT homologs are present in other annelids. We screened published transcriptomes and identified C_24_-SMTs homologs in three deep-sea gutless tubeworm species and in 17 gut-bearing annelid species from marine, limnic and terrestrial environments (**Supplementary Table 8**). Despite the presence of C_24_-SMT homologs in these annelids, the published sterol profiles of annelids, including four gut-bearing species analyzed in this study, are dominated by cholesterol (**Supplementary Table 8, Figure 1A)**. However, sitosterol and other methylated/ethylated sterols (C_28_ and C_29_ sterols) account for a considerable proportion of total sterols at 15-30% in some of these species, including the hydrothermal vent and cold seep tubeworms *Riftia pachyptila* and *Paraescarpia echinospica* (**Supplementary Table 8**). These deep-sea siboglinid annelids are only distantly related to *Olavius* and *Inanidrilus*, but also lack a gut and gain all of their nutrition from their chemosynthetic symbionts (Bright & Giere, 2005). The sterol contents of these tubeworms are dominated by cholesterol and desmosterol, but the C_28_ sterol campesterol and other phytosterols make up as much as nearly one third of their sterol contents (Guan et al., 2021; Phleger et al., 2005; Rieley et al., 1995). Our discovery of C_24_-SMTs homologs in these deep-sea annelids indicate that these tubeworms are able to synthesize phytosterols as well.

### Bona fide C_24_-SMT homologs are present in at least three animal phyla: sponges, rotifers and annelids

To assess the broader distribution of C_24_-SMT homologs among animals, we performed protein searches against public databases (see Materials and Methods for details). Hits were found in six other animal phyla: sponges, cnidarians, mollusks, nematodes, chordata and rotifers. Of these, the C_24_-SMT homologs detected in sponges, rotifers and annelids were identified as bona fide animal C_24_-SMTs (**Figure 3, Supplementary information**) and are described in the next paragraph. The C_24_-SMTs recovered from cnidarians, mollusks and chordata are unlikely to originate from the animals themselves. Their phylogenetic placement and high similarities to plant, algal and protist sequences suggest that these sequences originated from these animals’ diets or are contaminants (**Supplementary Figure 22 and Supplementary information**). We also found hits in some nematodes, but these belonged to a group of C-4 sterol methyltransferases (C_4_- SMTs), that are specific to nematodes (Chitwood, 1991; Darnet et al., 2020) and phylogenetically distinct from C_24_-SMTs (**Figure 3B, Supplementary Figure 22**).

**Figure 3.**
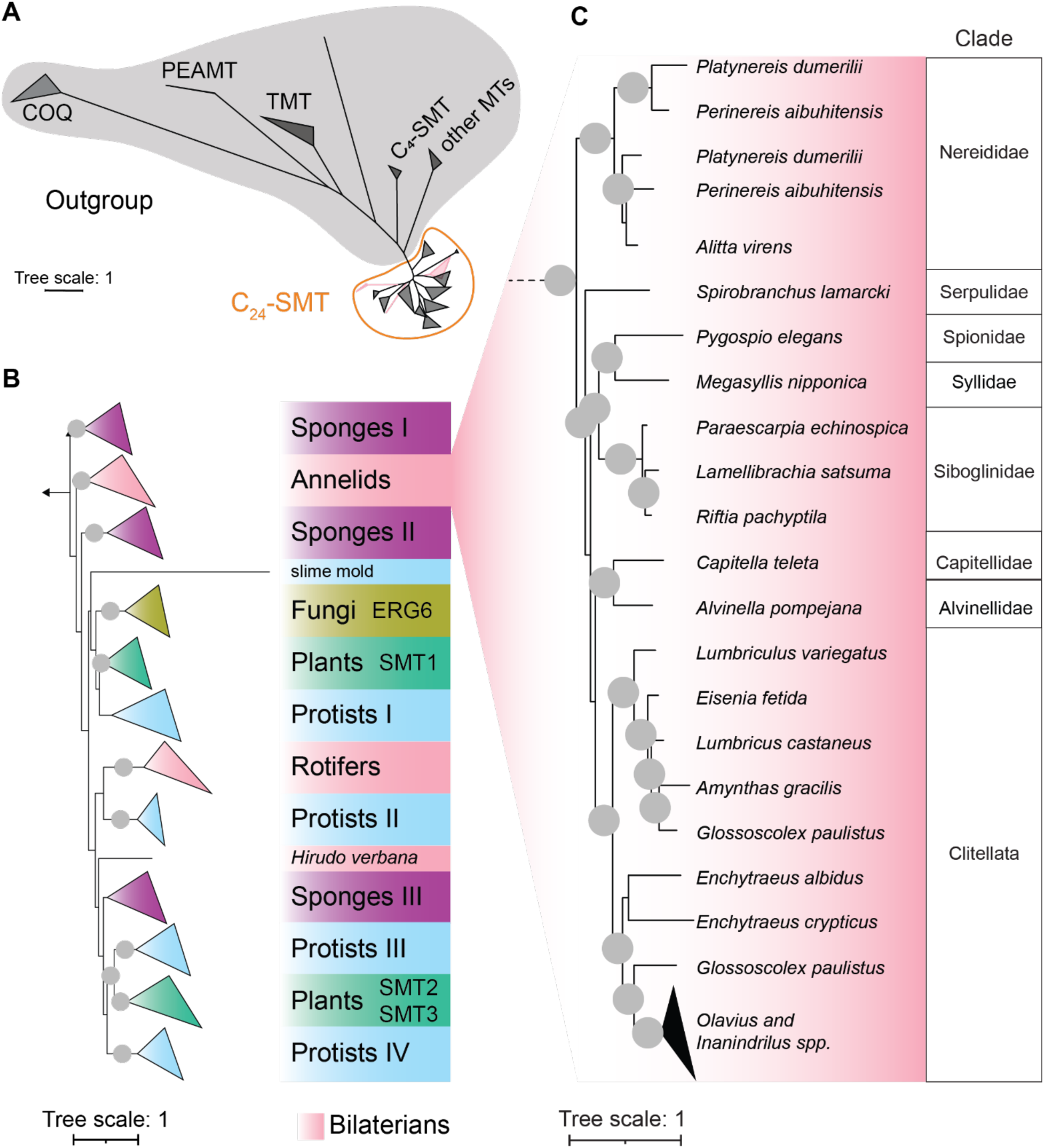
C_24_-SMTs are widely distributed across animals, particularly sponges, rotifers and annelids. **A**, The animal C_24_-SMT sequences cluster with bona fide plant and fungal C_24_-SMTs and are phylogenetically distant from other SAM-dependent methyltransferases (outgroup: Ubiquinone biosynthesis O-methyltransferase (COQ), phosphoethanolamine N-methyltransferase (PEAMT), tocopherol O-methyltransferase (TMT), C_4_ sterol methyltransferase (C_4_-SMT)). Unrooted maximum likelihood amino acid tree for eukaryotic SAM-dependent methyltransferases. **B**, Maximum likelihood amino acid tree of eukaryotic C_24_-SMTs. Sequences were clustered at 95% identity to make the tree more readable. Nodes with bootstrap value > 95 % are marked with grey circles. The tree is rooted at midpoint. **C**, C_24_-SMT homologs were detected in annelids from eight clades and were found in marine, limnic and terrestrial species.

Our phylogenetic analyses of bona fide C_24_-SMTs revealed that these fall into 12 clades (**Figure 3**). These 12 clades include canonical C_24_-SMTs from plants (SMT1 and SMT2/SMT3) and fungi (ERG6), which have been extensively characterized in previous studies (e.g. (Bouvier-Navé et al., 1998, p.; Haubrich et al., 2015; Nes et al., 2004)). Animal C_24_-SMTs do not cluster with these plant and fungi clades, suggesting that these canonical C_24_-SMTs were lost in the lineage leading to animals. Animal C_24_-SMTs fell in five subgroups. Three of these animal subgroups are from sponge C_24_-SMTs and are widely distributed across the tree. The remaining two animal clades contain sequences from annelids and rotifers (**Supplementary Information**) The annelid clade includes the C_24_-SMTs we discovered in gutless *Olavius* and *Inanidrilus*, and have shown are functional in producing sitosterol *de novo* in these animals. This clade also includes the C_24_-SMTs from all other annelids (26 species). These C_24_-SMTs are widely distributed across the annelid tree in eight orders, and in annelids from limnic, terrestrial and marine environments (**Figure 3**). Given that the phylogeny of C_24_-SMT corresponds well to the phylogenetic evolution of annelids (**Figure 3**), it is likely that C_24_-SMT was present in the last common ancestor of annelids.

## Conclusions

Here we describe and characterize a non-canonical animal C_24_-SMT that enables animals to synthesize sitosterol in a previously undescribed pathway that is distinct from that of plants.

Our phylogenetic analyses of C_24_-SMTs provide new insights into the evolutionary history of sterols in eukaryotes. While ubiquitous in plants and fungi, C_24_-SMTs were thought to be largely absent from animals (Haubrich et al., 2015; Volkman, 2005). Prior to this study, animal C_24_-SMT sequences had only been reported in a few sponges (one of the earliest diverging animal lineages) (Germer et al., 2017; Gold et al., 2016) and the annelid *Capitella teleta* (Najle et al., 2016), but the activity of these enzymes remains uncharacterized and their role in phytosterol synthesis was not explored. It has been assumed that C_24_-SMTs were present in the last eukaryotic common ancestor (LECA) and then lost in the animal branch (Desmond & Gribaldo, 2009; Gold et al., 2016; Haubrich et al., 2015). Our results, however, indicate that while the ancestral homologs that led to extant plant and fungal C_24_-SMTs were lost in the lineage leading to animals, they retained non-canonical C_24_-SMTs. These non-canonical C_24_-SMTs are present in at least three animal phyla, and sitosterol is the dominant sterol in at least some of these animals, particularly *Olavius* and *Inanidrilus*. Our analyses also revealed that the C_24_-SMTs sequences isolated from sponges and protists are widely distributed across the animal tree, suggesting that ancestral eukaryotes may have had several copies of the C_24_-SMT enzyme and flexible sterol synthesis pathways. While previous phylogenetic analyses indicated that at least one copy of the C_24_-SMT gene was present in LECA (Desmond & Gribaldo, 2009; Gold et al., 2016), our data suggest that more copies were present in the LECA and that the evolutionary history of sterol synthesis pathways is more complex than previously assumed. Our results furthermore challenge the widespread assumption that sterols can be used as geological biomarkers for teasing apart the evolutionary history of early plants and animals (Summons et al., 2022).

Given that C_24_-SMTs are widespread in animals, how can we explain the unusually high abundance of phytosterols in gutless *Olavius* and *Inanidrilus*? As described above, other gutless animals that gain their nutrition from chemosynthetic symbionts (such as vent and seep tubeworms) also have C_28_ and C_29_ sterol, but only *Olavius* and *Inanidrilus* have higher amount of sitosterol than cholesterol. Sterols play many essential roles in eukaryotes and their homeostasis is tightly regulated by complex mechanisms (Luo et al., 2020; Wollam & Antebi, 2011). Animal membranes are usually dominated by cholesterol, but studies have shown that phytosterols can be incorporated into animal membranes (Mouritsen & Zuckermann, 2004) and can have beneficial effects on animals. For example, they act as cholesterol-lowering agents, have anti-tumor, anti-inflammatory, antibacterial and antifungal properties (Bin Sayeed & Ameen, 2015; Saeidnia et al., 2014), and modulate interactions between bacterial pathogens and eukaryotic hosts (van der Meer-Janssen et al., 2010). Therefore, the anti-inflammatory and antibacterial properties of sitosterol, as well as its ability to protect animal cells against toxins that target cholesterol (Li et al., 2015), might play a role in the symbiosis between *Olavius* and *Inanidrilus* and their chemoautotrophic symbionts, by preventing the symbionts from entering the host cytoplasm. In addition, changes in sterol composition affect the fluidity and permeability of membranes, and these physical changes in turn affect many cellular processes. For example, the high levels of sitosterol in *Olavius* and *Inanidrilus* might increase the permeability of their membranes for dissolved gases from their environment. These hosts compete with their aerobic symbionts for the little oxygen available in their environment. Under oxic conditions, these worms’ symbionts consume 90% of the oxygen taken up by the worms (Häusler, Lott and Dubilier, unpublished results), indicating that the worms may often experience oxygen limitation. Furthermore, sitosterol has been shown to enhance mitochondrial energy metabolism in a mouse cell line (Shi et al., 2013), and might enable *Olavius and Inanidrilus* to gain more energy under low oxygen concentrations in their environment. While additional studies are needed to elucidate the physiological and/or ecological roles of sitosterol in animals, *Olavius and Inanidrilus* are valuable model systems for studying the impact of phytosterols on animal membrane properties *in vivo* and furthering our understanding of the roles sterols play in eukaryotic cells.

## Materials and Methods

### Reagents

All organic solvents were LC–MS grade: acetonitrile (ACN; Honeywell, Honeywell Specialty Chemicals), chloroform (Merck), isopropanol (IPA; BioSolve), methanol (MeOH; BioSolve), hexane (Sigma-Aldrich), acetone (Sigma-Aldrich), ethanol (EtOH; Sigma-Aldrich) and formic acid (FA; Sigma-Aldrich). Water was deionized using the Astacus MembraPure system (MembraPure). Pyridine (dried (max. 0.0075% H_2_O) SeccoSolv®) was obtained from Sigma-Aldrich. The reagents used for GC-MS derivatization were obtain from Chromatographie Service and Sigma-Aldrich. The internal standards (5α-cholestane and Ribitol) used for GC-MS analysis were obtained from Sigma-Aldrich.

### Sampling

#### Gutless annelids

Sediments in which gutless annelids occur were collected by scuba diving. The worms were extracted manually from the sediment and either directly fixed in MeOH or kept in aquaria with seagrass and sediment from the collection site for up to one year before use in experiments. Six gutless annelid species, belonging to two different genera were collected in five locations: the bay of Sant’Andrea (Island of Elba, Italy) (42° 48’29.4588” N; 10° 8’ 34.4436” E), the bay of Magaluf (Mallorca, Spain) (39º 30’ 14.814’’ N; 2º 32’ 35.868’’ E), at Carrie Bow Cay (Belize) (16° 04’ 59” N; 88° 04’ 55” W), Twin Cayes (Belize) (16° 50’ 3” N; 88° 6’ 23” W), and Okinawa (Japan) (26° 29’ 33.4” N; 127° 50’ 31.6” E).

#### Gut-bearing annelids

*Cirratulidae sp*., *Heronidrilus sp*. and *Rhyacodrilus sp*. were collected in Belize in the same environments as the gutless annelids. They were extracted manually from the sediment and directly fixed in MeOH. *Tubifex tubifex* specimens were purchased in an aquarium shop and kept in aquaria for two weeks before fixation in MeOH. *Capitella teleta* worms were provided by the Meyer Lab (https://wordpress.clarku.edu/nmeyer/) and kept in aquarium before fixation in MeOH. The MeOH fixed samples were stored at - 20°C until extraction.

#### Seagrass

Seagrass plants (*Posidonia oceanica*) were collected by scuba diving in the bay of Sant’Andrea (Elba, Italy) (42° 48’ 29.4588” N; 10° 8’ 34.4436” E). The leaves, roots and rhizomes were dissected using a razor blade, placed into individual bags and stored at -20°C.

#### Porewater

Porewater was collected from sediments for metabolomic analyses. We sampled in and near seagrass meadows in the Mediterranean Sea in the bay of Sant’Andrea (Elba, Italy) (42° 48’ 29.4588” N; 10° 8’ 34.4436” E) and in the Caribbean at Carrie Bow Cay (Belize) (16° 04’ 59” N; 88° 04’ 55” W) and Twin Cayes (Belize) (16° 50’ 3” N; 88° 6’ 23” W). Using a steel lance (1 m long, 2 mm inner diameter) fitted with a wire mesh (63 μm) to prevent the intake of sediment and seagrass, porewater was slowly extracted from sediments into polypropylene syringes. A porewater profile consisted of top to bottom sampling of the sediments every 5 or 10 cm down to 30 cm. For metabolomic analysis, 10 mL samples were stored at -20°C until further processing.

### Metabolite extraction

#### Gutless and gut-bearing annelids

Metabolites were extracted from the worms using the following method for metabolite profiling: Tissues from MeOH fixed worms were transferred to 2 mL screwcap tubes containing a mix of silica beads (Sigmund Linder). The residual methanol was added to the screw cap tube. The tubes were spiked with 100 μL 5α-cholestane (1 mM) and 40 μL ribitol (0.2 mg mL^-1^). 0.5 mL pre-cooled extraction solution (ACN:MeOH:water (v:v:v) 2:2:1) was added to each tube.

Tissues were disrupted by bead beating for 2 cycles of 40 sec (4 m sec^-1^). The tissues were pelleted by centrifugation (9,600 x g, 2 min), and the supernatants transferred to new tubes. The pellets were extracted one more time with 1.5 mL of extraction solution. The supernatants were combined and evaporated to dryness in a vacuum concentrator without heating (approximately 1.5 h). The obtained aliquots were stored at -20°C until metabolite derivatization.

#### Seagrass

The frozen plant tissues were ground to a fine powder in liquid nitrogen using a pestle and mortar. 70 mg of the powder was transferred to 2 mL screw cap tubes containing 1.2 mL MeOH (pre-cooled at -20°C). The tubes were vortexed for 10 s. The internal standards were added to the tubes: 40 μL ribitol (0.2 mg mL^-1^) and 100 μL 5α-cholestane (1 mM), and the tubes were vortexed for another 10 sec. The tubes were placed on a thermomixer and shaken for 10 min at 70°C at 950 rpm. The plant powder was pelleted by centrifugation (10 min, 11,000 × g, 4°C). The supernatant was transferred into a new 2 mL Eppendorf tube and evaporated to dryness using a Concentrator Plus (Eppendorf) (V-AL, 1.5 h, 45°C). The dried extracts were stored at - 20°C until metabolite derivatization.

#### Porewater

Sterols were extracted using Superclean LC-18 SPE tubes (6 mL, 0.5 g, Supelco). The silica cartridge was equilibrated using an ultra-pure water (UPW):MeOH dilution series (0:1, 1:4, 1:1, 4:1, 1:0 [v/v]). The porewater samples were spiked with internal standard (100 μL 5α-cholestane (1mM)), before being loaded on the column. Impurities were removed by three successive UPW washes, and the sterols eluted from the cartridge with 3 × 5 mL of MeOH. The MeOH fractions were collected and evaporated to dryness using a Concentrator Plus on V-AL mode with centrifugation at 30°C. Positive and negative controls were run in parallel. For negative controls sterols were extracted from 10 mL of artificial sea water (ASW). As with positive controls, 10 mL ASW was spiked with 20, 40, or 80 nmol cholesterol and β-sitosterol. The dried extracts were stored at -20°C until metabolite derivatization.

### GC-MS analysis

#### Derivatization

To remove condensation formed during storage, we further dried the extracts in a vacuum concentrator for 30 min prior to sample preparation for GC-MS analysis.

#### Gutless and gut-bearing annelids

Metabolite derivatization was performed by adding 80 μL methoxyamine hydrochloride (MeOX) dissolved in pyridine (20 mg mL^-1^) to the dried pellet and incubating for 90 min at 37°C using a thermomixer (BioShake iQ, Analytik Jena) under constant rotation at 1350 rpm. Following the addition of 100 μL N,O-Bis(trimethylsilyl)trifluoroacetamide (BSTFA), each extract was vortexed, and incubated for another 30 min at 37°C on a thermomixer under constant rotation at 1350 rpm. After a short centrifugation, 100 μL of the supernatant was transferred to a GC-MS vial (Insert G27, sping S27 and Mikor-KH-Vial G1; Chromatographic Service) for GC-MS data acquisition.

#### Seagrass

80 μL of MeOX (20 mg mL^-1^ dissolved in pyridine) was added to the dried extracts. The resuspended dried extracts were vortexed for a few seconds and placed on a thermomixer (BioShake iQ, Analytik Jena) for 90 min (37°C, 1200 rpm). 80 μL BSTFA was added to the tubes. The tubes were vortexed for a few seconds and placed on a thermomixer for 15 min (60°C, 1200 rpm). After a short centrifugation, 100 μL of the supernatant was transferred into GC-MS vials (Insert G27, sping S27 and Mikor-KH-Vial G1; Chromatographic Service) and analyzed by GC-MS.

#### Porewater

After complete evaporation, 80 μL BSTFA was added to the tubes. The tubes were gently vortexed and placed on a thermomixer for 15 min (60°C, 950 rpm). After a short centrifugation (1 min, 7,800 x g), the supernatant was transferred into GC-MS vials and analyzed.

#### Data acquisition

The analysis of all metabolomic samples was conducted on a 7890B GC system (Agilent Technologies) coupled to a 5977A single quadrupole mass selective detector (Agilent Technologies). The gas chromatograph was equipped with a DB-5 ms column (30 m × 0.25 mm, film thickness 0.25 μm; including 10 m DuraGuard column, Agilent Technologies) and a GC inlet liner (ultra inert, splitless, single taper, glass wool, Agilent). Helium was used as gas carrier at a constant flow (0.8 mL min^-1^). An Agilent 7693 autosampler injected 1 μL of derivatized sample in splitless mode. The injector temperature was set at 290°C. The temperature program started at 60°C for 2 min, then increased to 300°C at 10°C min^-1^, and held at 325°C for 7 min. Mass spectra were acquired in electron ionization mode at 70 eV across the mass range of 50–600 *m/z* and a scan rate of 2 scans sec^-1^. The retention time was locked using a standard mixture of fatty acid methyl esters (Sigma-Aldrich).

#### Data analysis

Sterols were identified through comparison with standards using the Mass Hunter Suite (Agilent) and through comparison to the NIST database. Sterols were further quantified using the Mass Hunter Quantification Suite (Agilent).

### HPLC-MS

#### High-resolution LC–MS/MS

The analysis was performed using a QExactive Plus Orbitrap (Thermo Fisher Scientific) equipped with a HESI probe and a Vanquish Horizon UHPLC system (Thermo Fisher Scientific). The lipids were separated on an Accucore C30 column (150 × 2.1 mm, 2.6 μm, Thermo Fisher Scientific), at 40°C, using a solvent gradient. Buffer A (60:40 ACN:H2O, 10 mM ammonium formate, 0.1% FA) and buffer B (90:10 IPA:ACN, 10 mM ammonium formate, 0.1% FA) (Breitkopf et al., 2017) were used at a flow rate of 350 μl min^−1^. The lipids were eluted from the column with a gradient starting at 0% buffer B (**Supplementary Table 10**). The injection volume was 10 μl. In the same run, MS measurements were acquired in positive-ion and negative-ion mode for a mass detection range of *m/z* = 150–1,500. Resolution of the mass analyzer was set to 70,000 for MS scans and 35,000 for MS/MS scans at *m/z* = 200. MS/MS scans of the eight most abundant precursor ions were acquired in positive-ion and negative-ion modes. Dynamic exclusion was enabled for 30 sec and collision energy was set to 30 eV (for more details see **Supplementary Table 11**). The data were analyzed with Thermo FreeStyle™ (version 1.6) and Xcalibur Quan Browser Software v. 2.0.3 (Thermo).

### GC-IRMS

#### Sample preparation

Sterols were extracted from the gutless annelids *O. algarvensis* and *O. ilvae*, as well as from the rhizome and leaves of *P. oceanica*. All samples were extracted with dichloromethane:MeOH (2:1) three times. The resulting total lipid extracts were then separated by solid phase extraction with a Machery & Nagel aminopropyl modified silica gel column (500mg) into four fractions with increasing polarity (see (Birgel et al., 2008)). For the sterols, the third fraction was used (dichloromethane:acetone 9:1). Prior to measurement on the GC-MS and GC-IRMS, the samples were silylated with BSTFA. The porewater samples were extracted as described above.

#### Data acquisition

The resulting sterols were identified on a Thermo Electron Trace DSQ II coupled gas chromatograph mass spectrometer (GC-MS). The GC-MS was equipped with a 30 m HP-5 MS UI fused silica capillary column (0.25 mm diameter, 0.25 μm film thickness). The carrier gas was helium. The GC temperature program was as follows: 60°C (1 min), from 60°C to 150°C at 10°C min^-1^, from 150°C to 325°C at 4°C min^-1^, 25 min isothermal. Identification of compounds was based on retention times and published mass spectral data. Compound-specific carbon stable isotope compositions of sterols were measured on a gas chromatograph (Agilent 6890) coupled with a Thermo Finnigan Combustion III interface to a Finnigan Delta Plus XL isotope ratio mass spectrometer (GC-IRMS). The GC conditions were identical to those described above for GC-MS analyses. All sterols were corrected for the additional carbons introduced by derivatization with BSTFA. The standard deviation of the isotope measurements was < 0.8‰.

### MALDI-2-MSI

#### Sample preparation

The worms were prepared following Kadesch et al. (2019) with a few modifications. Briefly, 20 μL 6.7% glutaraldehyde solution in marine phosphate buffered saline were deposited on a glass slide. Gutless annelids were transferred to the fixative using bent acupuncture needles. A coverslip was applied, and samples were frozen in liquid nitrogen and stored at −80°C until further processing.

A single worm was transferred using featherweight forceps onto a sodium carboxymethyl cellulose (CMC; Sigma-Aldrich) stamp which was glued onto a cryostat specimen disc (Leica). Gelatin solution (8 % (β = 80 g L^-1^), 10-20 μL) was used to coat the worm. The specimen disc with the sample was transferred to the cryostat (Leica CM3050 S, Leica Biosystems) and kept for 30 min at -22°C before sectioning into sections of 12 μm thickness (chamber temperature -22°C, object temperature -22°C). Sections were thaw-mounted on IntelliSlides (Bruker Daltonics), and their quality determined microscopically. Sections of sufficient quality were stored in a desiccator at room temperature (RT) until further analysis.

#### Matrix application and data acquisition

The matrix 2′,5′-dihydroxyacetophenone (DHAP; Merck) was applied by sublimation and the data were acquired on a modified timsTOF fleX instrument (Bruker Daltonics, Bremen, Germany) (Soltwisch et al., 2020) as described in (Bien et al., 2021). The mass resolving power of this hybrid QTOF-type instrument is about 40,000 (fwhm) in the investigated *m/z* range of 300–1500. For material ejection, a scan range of 1 μm of the laser spot on the target was used, resulting in an ablated area of 5 μm in diameter. The step size of the stage during the MSI run was set to 5 μm. The laser power was set to 40%, with 50 laser shots/pixel.

#### Data analysis

SCiLS lab (Bruker Daltonics, version 2021a) was used for data analysis and to produce the ion images shown in the figures. For image visualization, an interval width of 15 ppm was used. The ion images represent the data after root mean square normalization and without denoising.

### TOF-SIMS

#### Sample preparation

*O. algarvensis* specimens were fixed with 4% paraformaldehyde (PFA) at 4°C for 4 h. The fixative was removed by washing three times with marine phosphate buffer. The washed samples were then stored at -20°C in MeOH.

PFA-fixed samples were embedded in paraffin. The MeOH was exchanged with pure EtOH by three successive incubations of 60 min in pure EtOH at RT. The samples were then incubated in RothiHistol (30 min, 60 min, and overnight at RT). The samples were then infiltrated with paraffin at 60°C, they were placed in fresh paraffin three times for 30, 60 and 60 min and then left to incubate overnight. For embedding, two-thirds of the embedding mold was filled with paraffin. The sample was placed in the mold and the mold was filled completely with paraffin. The sample was aligned and left to polymerize for one week. After polymerization, a microtome was used to cut 4 μm sections. The sections were placed on poly-L-lysine-coated glass slides (Sigma-Aldrich), left to air dry overnight and baked at 60°C for 2 h to improve adherence to the slide. Finally, the sections were de-waxed, first in three baths of 10 min in RothiHistol, followed by an EtOH series (96%, 80%, 70%, 50%), and the slides were dipped in ultra-pure water and left to air dry. Once dried they were wrapped in aluminum foil and stored in a desiccator (Roth, Desiccator ROTILABO® Glass, DN 250, 8.0 l) until TOF-SIMS analysis.

#### Data acquisition and analysis

SIMS data were acquired on an IONTOF TOF-SIMS 5 instrument (IONTOF GmbH) using a 25 keV Bi_3_^+^ LMIG analytical beam in positive mode. To obtain high-resolution mass spectra, high current bunch mode was used with a beam current of ∼1 pA. The analytical area was typically 100 μm^2^. Secondary ion maps were collected using the Bi_3_^+^ LMIG in burst alignment mode for greater lateral resolving power (raster size: 512 × 512, FoV 210 × 210). The samples were also sputter pre-cleaned using a 5 keV Ar_1000_^+^ cluster ion beam. The data were analyzed in SurfaceLab 7 (IONTOF). We analyzed standards to determine the most abundant ions produced by each sterol: cholesterol ([M-H_2_O+H]^+^ C_27_H_45_, at *m/z* 369.38; [M-H]^-^ C_27_H_45_O, at *m/z* 385.34; [M-H+O]^-^ C_27_H_45_O_2_, at *m/z* 401.35) and sitosterol ([M-H_2_O+H]^+^ C_29_H_49_, at *m/z* 397.47; [M-CH_3_O]^-^ C_28_H_47_, at *m/z* 383.37; [M-H]^-^ C_29_H_49_O, at *m/z* 413.45). To determine the distribution of each sterol in *O. algarvensis* sections, the intensity of the three most abundant ions was combined.

### Sterol identification

Sterols were identified by comparison to chemical standards and MS database. Matching mass spectra and retention time with sterol standards confirmed sterol identification. In addition, tandem mass spectra were acquired with high-mass resolution and accuracy on all sterols. Each sterol was identified on using a combination of two different chromatography types (GC-MS and LC-MS), including different ionization methods (electrospray ionization, electron impact ionization). For sterol identification with MSI we used chemical standards for sterols and measured them with MALDI-2 and SIMS in parallel to the tissue sections

### Identification of genes involved in sterol biosynthesis in *O. algarvensis* transcriptomes

Transcriptomes generated in a previous study (Wippler et al., 2016) were analyzed for genes involved in sterol biosynthesis Protein sequences from humans, *Arabidopsis thaliana* and *Saccharomyces cerevisiae* were used as queries (**Supplementary Table 12**) to search the transcriptomic assemblies with TBLASTN (e-value 1e-10). The identity of the hits was confirmed by BLASTP search against the NCBI nr and Swiss-Prot database as well as by INTERPROSCAN domain prediction. *O. algarvensis* sequences were aligned with reference sequences (Desmond & Gribaldo, 2009) using Clustalw (Larkin et al., 2007), trimmed with trimAI (Capella-Gutiérrez et al., 2009). Alignments were used to calculate maximum-likelihood trees with ultrafast bootstrap support values using IQ-TREE (Minh et al., 2020). The resulting trees were visualized using iTOL (Letunic & Bork, 2019). The trees are shown in **Figure 3** and **Supplementary Figures 5** to **14**.

### Gutless annelid nucleic acid extraction, sequencing and analysis

#### Extraction and sequencing

Genomic DNA was extracted from fresh specimens of two gutless annelid species (*O. algarvensis* and *O. ilvae*, one individual each). High molecular weight genomic DNA was isolated with the MagAttract HMW DNA Kit (Qiagen). Quality was assessed by the Agilent FEMTOpulse and DNA quantified by the Quantus dsDNA kit (Promega). DNA was processed to obtain a PacBio Sequencing-compatible library following the recommendations outlined in “Procedure & Checklist – Preparing HiFi Libraries from Low DNA Input Using SMRTbell Express Template Prep Kit 2.0”. Libraries were sequenced on a Sequel II instrument at the Max-Planck Genome-Centre Cologne (MP-GC) with sequencing chemistry 2.0, binding kit 2.0 on one 8M SMRT cell for 30 h applying continuous long read (CLR) sequencing mode.

#### Assembly and identification of genes involved in sterol biosynthesis

The CLR reads were assembled using Flye (v 2.8) (Kolmogorov et al., 2019). The completeness of the assembly was assessed with QUAST (v. 5.0.0) (Gurevich et al., 2013) and BUSCO (version 5.2.2) (Seppey et al., 2019). *O. algarvensis* and *O. ilvae* sequences were retrieved from the PacBio assembly with BLAT (Kent, 2002) and SCIPIO (version 1.4) (Keller et al., 2008) using *O. algarvensis* transcripts as queries.

### Metaproteomics

#### Protein identification and quantification

We re-analyzed data produced by Jensen (Jensen et al., 2021) using a customized database containing 1,439,433 protein sequences including host and symbiont proteins as well as a cRAP protein sequence database (http://www.thegpm.org/crap/) of common laboratory contaminants. We performed searches of the MS/MS spectra against this database with the Sequest HT node in Proteome Discoverer version 2.3.0.523 (Thermo Fisher Scientific). The following parameters were used: trypsin (full), maximum two missed cleavages, 10 ppm precursor mass tolerance, 0.1 Da fragment mass tolerance and maximum of 3 equal dynamic modifications per peptide, namely: oxidation on M (+ 15.995 Da), carbamidomethyl on C (+ 57.021 Da) and acetyl on the protein N terminus (+ 42.011 Da). False discovery rates (FDRs) for peptide spectral matches (PSMs) were calculated and filtered using the Percolator Node in Proteome Discoverer (Spivak et al., 2009). Percolator was run with a maximum delta Cn 0.05, a strict target FDR of 0.01, a relaxed target FDR of 0.05 and validation based on q-value. We used the Protein FDR Validator Node in Proteome Discoverer to calculate q-values for inferred proteins based on the results from a search against a target-decoy database. Proteins with a q-value <0.01 were categorized as high-confidence identifications and proteins with a q-value of 0.01– 0.05 were categorized as medium-confidence identifications. We combined search results for all samples into a multi-consensus report in Proteome Discoverer and only proteins identified with medium or high confidence were retained, resulting in an overall protein-level FDR of 5%.

### C_24_-SMT distribution in animals

#### Gutless annelid search

Total RNA was extracted from nine gutless annelid species. RNA was quality and quantity assessed by capillary electrophoresis (Agilent Bioanalyser PicoChip) and Illumina-compatible libraries were generated with the NEBNext® Single Cell/Low Input RNA Library Prep Kit for Illumina®. Libraries were again quality and quantity controlled followed by sequencing on a HiSeq 3000 device with 2 × 150 bp paired-end read mode. The raw reads were trimmed and corrected, the symbiont reads were mapped out and the host read assembled using Trinity. The resulting assemblies were screened for C_24_-SMT homologs using the model C_24_-SMTs (P25087, Q9LM02, Q39227) as subject. Only hits with e-values smaller than 1e-30 and spanning at least half the subject sequences were kept for further analysis. The contigs of interest were isolated and TransDecoder (v 5.5.0) (https://github.com/TransDecoder) was used to identified candidate coding region TransDecoder, homology searches (blastp and PFAM search) were used as additional retention criteria.

#### Search in public databases

To assess the presence of C_24_-SMT homologs in other animals, protein searches against NCBI (National Center for Biotechnology Information (NCBI), available from: https://www.ncbi.nlm.nih.gov/) databases (nr, tsa_nr, refseq_prot, env_nr and tsa) as well as the proteomes predicted by Ensembl metazoan (Howe et al., 2020) and Compagen (Hemmrich & Bosch, 2008) were performed. The search was restricted to metazoan sequences to avoid hits from fungi and viridiplantae. Only hits with e-values smaller than 1e-30 and a coverage > 60% were retained for further analysis.

#### Phylogenetic tree

A phylogenetic tree was constructed from selected C_24_-SMT protein sequences. Briefly, published C_24_-SMT sequences (Desmond & Gribaldo, 2009; Gold et al., 2016) were downloaded from UniProt and JGI Genome Portal and used as references. Other SAM-dependent methyltransferases (ubiquinone biosynthesis O-methyltransferase, phosphoethanolamine N-methyltransferase, tocopherol O-methyltransferase and C_4_-sterol methyltransferase) were used as outgroups. Together with the animal C_24_-SMTs these sequences were clustered (95% ID), aligned using Clustalw 2.1 (Larkin et al., 2007) and trimmed with TrimAI v1.2 (Capella-Gutiérrez et al., 2009). IQ-TREE (v1.6.12) (Minh et al., 2020) was used to predict the best-fit models of evolution and to calculate a maximum-likelihood tree with ultrafast bootstrap support values from the concatenated alignment. The resulting tree was visualized using iTOL v6 (Letunic & Bork, 2019).

### C_24_-SMT heterologous gene expression and enzyme assay

#### Heterologous gene expression

To determine the activity of the putative C_24_-SMT enzymes, we overexpressed OalgSMT, OclaSMT and ArathSMT1 (**Supplementary Table 13**) in *E. coli* and performed enzymatic assays. GenScript (Genscript®) generated pet28a(+) (NheI/XhoI) plasmid containing the sequence of interest. For expression, the pet28(a)-OalgSMT and pet28(a)-OclaSMT vectors were transformed in Lemo21(DE3) *E. coli* competent cells (New England Biolabs (NEB)). The pet28(a)-ArathSMT1 vector was transformed in Overexpress C43(DE3) pLysS *E. coli* competent cells (Lucigen). A single colony of transformed cells was grown in 3 mL lysogeny broth (LB) supplemented with the appropriate antibiotics for 8 h (37°C, 150 rpm). 1 mL pre-culture was used to inoculate 1 L ZYP-5052-Rich-Autoinduction-Medium (Studier, 2005) supplemented with antibiotics and 1500 μM rhamnose. The cultures were grown for 72 h at 20°C, with rotation at 150 rpm. Cells were harvested by centrifugation at 4,500 × g for 25 min at 4°C, the supernatant discarded and the resulting pellet stored at -20°C until further use.

#### Protein extraction/cell lysis

The frozen pellets were thawed on ice. They were then resuspended in 15 mL sucrose solution (750 mM sucrose solution in 20 mM phosphate buffer at pH 7.5). Once the pellets were dissolved in the sucrose solution, 5 mg lysozyme was added and the tubes shaken for 10 min at RT. Cells were lysed by addition of 30 mL lysis buffer (100 mM NaCl, 15% glycerol, 3 mM EDTA in 20 mM phosphate buffer at pH 7.5), 0.2 mL MgSO_4_ stock solution (120 g L^-1^) and 0.3 mL Triton X-100. The mix was vigorously shaken and incubated on ice until it reached a gelatinous consistency. The DNA was fragmented by addition of 2 mL DNase stock solution (50 mg DNase I in 35 mL buffer (20 mM Tris-HCl pH8, 0.5 M NaCl) and 15 mL glycerol). Finally, the cell fragments and inclusion bodies were pelleted by centrifugation (45min, 16,000 × g, 4°C). The supernatant, containing soluble proteins, was aliquoted and stored at -80°C until further use. The overexpression yielded a target protein migrating on SDS-PAGE with the expected size of ∼40 kDa.

#### Enzymatic assay

We tested nine sterol substrates (cycloartenol, lanosterol, zymosterol, lathosterol, 7-dehydrocholesterol, desmosterol, campesterol, cholesterol and 24-methylene-cholesterol) (**Supplementary Table 14**). The assay was performed in 600 μL total volume. 100 μL crude soluble protein extract was mixed with 400 μL phosphate buffer (20 mM, 5% glycerol, pH 7.5) in a 15 mL tube containing a sterol substrate (final concentration 100 μM) dispersed in Tween 80 (0.1% v/v). The reaction was initiated with 100 μL SAM working solution (0.6 mM). The reaction was performed in a water bath at 35°C for 16 h. The reaction was terminated with 600 μL 10% methanolic KOH. The products were extracted three times with 2.5 mL hexane and mixed on a vortex for 30 s. The resulting organic layer was evaporated to dryness in a concentrator at 30°C, V-AL for 1.5 h. Two internal standards, 5α-cholestane (100 μL, 1 mM solution) and ribitol (40 μL 200 mg/L solution), were added to the tubes and evaporated to dryness. The samples were derivatized and analyzed on an Agilent GC-MS as described above or directly analyzed on a QExactive Plus Orbitrap (Thermo Fisher Scientific) equipped with a HESI probe and a Vanquish Horizon UHPLC system (Thermo Fisher Scientific). The sterols were separated on a C18 column (Accucore Vanquish C18+, 100 × 2.1, 1.5 μm, Thermo Fischer Scientific), for method details see **Supplementary Table 11 and 15**.

## Data availability

The metaproteomic mass spectrometry data have been deposited at the ProteomeXchange Consortium via the PRIDE partner repository (Vizcaíno et al., 2016) with the following dataset identifiers: PXD014881.

The upload of metabolomics and sequencing data on public platforms is in progress.

### Author Contributions

D.M., M.L. and N.D. conceived the study. D.M. collected, processed and analyzed the metabolomic, metatranscriptomic and metagenomic data. T.B. and S.F. collected mass spectrometry imaging data. D.B. performed the GC-IRMS measurements. M.J. and M.K. collected and analyzed the proteomics data. C.Z. and D.M. performed the heterologous gene expression. D.M. performed the enzyme assay and analyzed the data. D.M. and M.L. wrote the manuscript together with N.D. and contributions from T.B., D.B., M.K., and C.Z..

## Acknowledgments

We thank Alfred Garsdal, Janine Beckmann, Marvin Weinhold, Martina Meyer, Silke Wetzel, Tomasz Markowski, Tora Gulstad, Kristopher Caspersen and Frantisek Fojt from the Max Planck Institute for Marine Microbiology (MPI-MM) for support with data acquisition and sample preparation. We thank Miriam Sadowski and Grace D’Angelo for proof reading the manuscript. We are grateful to Alexander Gruhl, Yui Sato, Niko Leisch, Anna Mankowski, Silvia Bulgheresi, Claudia Bergin, Emilia M. Sogin and Katherine Schimdt for sample collections and help in the field. We thank the Meyer Lab for providing us with *Capitella teleta* worms. We also thank Bruno Huettel (Max Planck Genome Center, Cologne) for his support with sequencing. LC-MS/MS based proteomics measurements were made in the Molecular Education, Technology, and Research Innovation Center (METRIC) at North Carolina State University. We thank Bruker Daltonics for providing access to the prototype timsTOF flex MALDI-2 instrument and Jen Soltwisch and Klaus Dreisewerd for their advice and support during the sterols imaging. We are grateful for fruitful discussions with Liz Hambleton, with Christopher Laumer and with colleagues in the Department of Symbiosis (MPI-MM). We thank Miriam Weber, Christian Lott, and HYDRA staff for sample collections. We are grateful to the Max-Planck Society for their generous financial support over the many years it took to bring this work to fruition. M.K. acknowledges financial support from the National Institute of General Medical Sciences of the National Institutes of Health under Award Number R35GM138362 and the U.S. National Science Foundation (grant IOS 2003107). This work is contribution *XXX* from the Carrie Bow Cay Laboratory, Caribbean Coral Reef Ecosystem Program, National Museum of Natural History, Washington DC.

