## Supplementary material for "*De novo* phytosterol synthesis in animals"

### Table of content

Supplementary Information ............ 2-4

Supplementary Figures ............ 5-26

Supplementary Tables ............ 27-36

References ............ 37-40

### Supplementary Information

**Sitosterol is present in the environment of the gutless annelids in concentrations sufficient to sustain the growth of small sterol-auxotrophic invertebrates.** Chemical analysis of pore water profiles collected in the vicinity of seagrass meadows, the habitat of many gutless annelids, revealed an irregular distribution of sterols. Some samples, such as those collected near *Posidonia oceanica* seagrass meadows off the island of Elba (Mediterranean), had cholesterol and sitosterol present in the nano- to micro-molar range **(Supplementary Figure 3).** Samples collected in Belize (Caribbean), in the vicinity of the seagrasses *Thalassia testudinum* and *Syringodium filiforme,* were devoid of detectable amounts of sitosterol or cholesterol. Seagrasses, like terrestrial plants, exude organic compounds into the substrate surrounding their roots, the rhizosphere (Sogin et al., 2022; Vives-Peris et al., 2020). The sterol profile of *P. oceanica* roots was composed of sitosterol (69%), stigmasterol (11%) and campesterol (20%). *P. oceanica* could thus be the origin of the sitosterol in the porewater, but cannot be the source of cholesterol as it was not present in its tissues. Cholesterol and sitosterol concentrations measured in the porewater environment (ranging from 25 nM to 3 μM) are in the range of reported minimal dietary sterol requirements for small sterol-auxotrophic invertebrates (Carvalho et al., 2010; Lu et al., 1977). Furthermore, the sterols in the porewater could pass through the cuticle of the worm as it is permeable to substances up to 70 kDa (Dubilier et al., 2006).

**The isotopic signature of the sterols in the worms exclude an environmental origin**. Results from GC-IRMS with single metabolite resolution showed that the sitosterol present in seagrass tissue had $\delta^{13}C$ values ranging from −21 ‰ to −15 ‰ (**Figure 1E**).

These values are in accordance with bulk measures of isotopic composition for *P. oceanica* which range from −16.4 ‰ to −8.3 ‰ (Cooper & DeNiro, 1989; Jennings et al., 1997; Lepoint et al., 2004; McMillan et al., 1980; Pinnegar & Polunin, 2000; Vizzini et al., 2010), and also match values previously reported for sterols in other seagrasses (Canuel et al., 1997). Sterols from sediment porewater had $\delta^{13}C$ values of −30 ‰ to −26 ‰ (**Figure 1E**), which are similar to previously reported ranges (Canuel et al., 1997; Dauby, 1989; Fry et al., 1983; Thayer et al., 1978) and about 10 ‰ lower than the sterols from *P. oceanica*. This difference reflects the mixed sources of the sterols present in the sediment, most of which are likely planktonic in origin.

**The total sterol content of gutless annelids is comparable to the content of other worms.** Measurements of metabolites in single *Olavius algarvensis* worms with mass spectrometry revealed a total free sterol content of 3.44 ± 0.04 µg ($n$ = 12). Assuming an average wet weight per worm of 1 mg, sterols represent 0.34 % wet weight, a value similar to reported percentages for terrestrial and aquatic worms (Ballantine et al., 1978; McLaughlin, 1971; Voogt, 1973a; Wilber & Bayors, 1947).

**SMTs in rotifers**. $C_{24}$-SMT homologues were detected in nine rotifers species. The rotifer sequences grouped together to form a sister group to unicellular eukaryotes SMT (**Figure 3, Supplementary Figure 23**). Presence of $C_{24}$-SMT homologues in 2 genera and the fact that the rotifers were isolated from different environments and had access to different food resources, suggests that most recovered sequences were not a result of contamination. The only exception are the sequences isolated from the two *Branchionus* species, which do not cluster with the other rotifers sequences but with algae and protists sequences.

Our results suggest that different SMT isoforms are present in rotifers. We identified up to four different SMT copies in a single species (**Supplementary Figure 23**). Sterol auxotrophy has been proposed for rotifers (Wacker & Martin-Creuzburg, 2012) and $C_{24}$-SMTs might play a different role in these organisms. Animal sterol auxotrophs usually lack the three enzymes responsible for the transformation of squalene into lanosterol: farnesyl-diphosphate farnesyltransferase 1 (FDFT1), squalene monooxygenase (SQE) and lanosterol synthase (LAS) (Shamsuzzama et al., 2020). However, the presence of downstream cholesterol synthesis genes has been reported in sterol auxotrophs. Some of these sterol enzymes retain their catalytic functions and are involved in the conversion of dietary sterols into cholesterol (Meyer et al., 1979; Shamsuzzama et al., 2020), while others now have distinct functions (Oh et al., 2017; Shamsuzzama et al., 2020). The function performed by $C_{24}$-SMT enzymes in rotifers would benefit from further study.

**$C_{24}$-SMT homologues detected in cnidarian, chordata and mollusks are likely to be contaminations.** Putative $C_{24}$-SMT homologs that were phylogenetically very distant to the bona fide animal $C_{24}$-SMTs found in annelids, sponges and rotifers were found in the transcriptomes of animals from three other animal phyla: cnidarian, chordata and mollusks (**Figure 3, Supplementary Figure 22**). The cnidarian sequence cluster with the Sponge II group, in absence of more cnidarian sequences it is not possible to determine if this sequence is $C_{24}$-SMT homolog present in cnidarians or a sponge contamination. We found a unique sequence in the chordata phyla. This chordata sequence closely resembles the $C_{24}$-SMT of *Oryza sativa* and clusters with the plant SMT2/SMT3 sequences and was therefore considered as a plant contamination. Several sequences were isolated from mollusks species. However, given their broad and discontinuous taxonomic affiliations and phylogenetic distance to the sponge, annelid and rotifer $C_{24}$-SMTs, these homologs may not be bona fide animal $C_{24}$-SMTs. Their high similarities to plant, algal and protist sequences, and the phylogenetic distances between putative $C_{24}$-SMTs isolated from species belonging the same genus (*Haliotis*) (**Supplementary Figure 22**) led us to conclude that these sequences likely originated from these animals' diets or are contaminants.

### Supplementary Figures

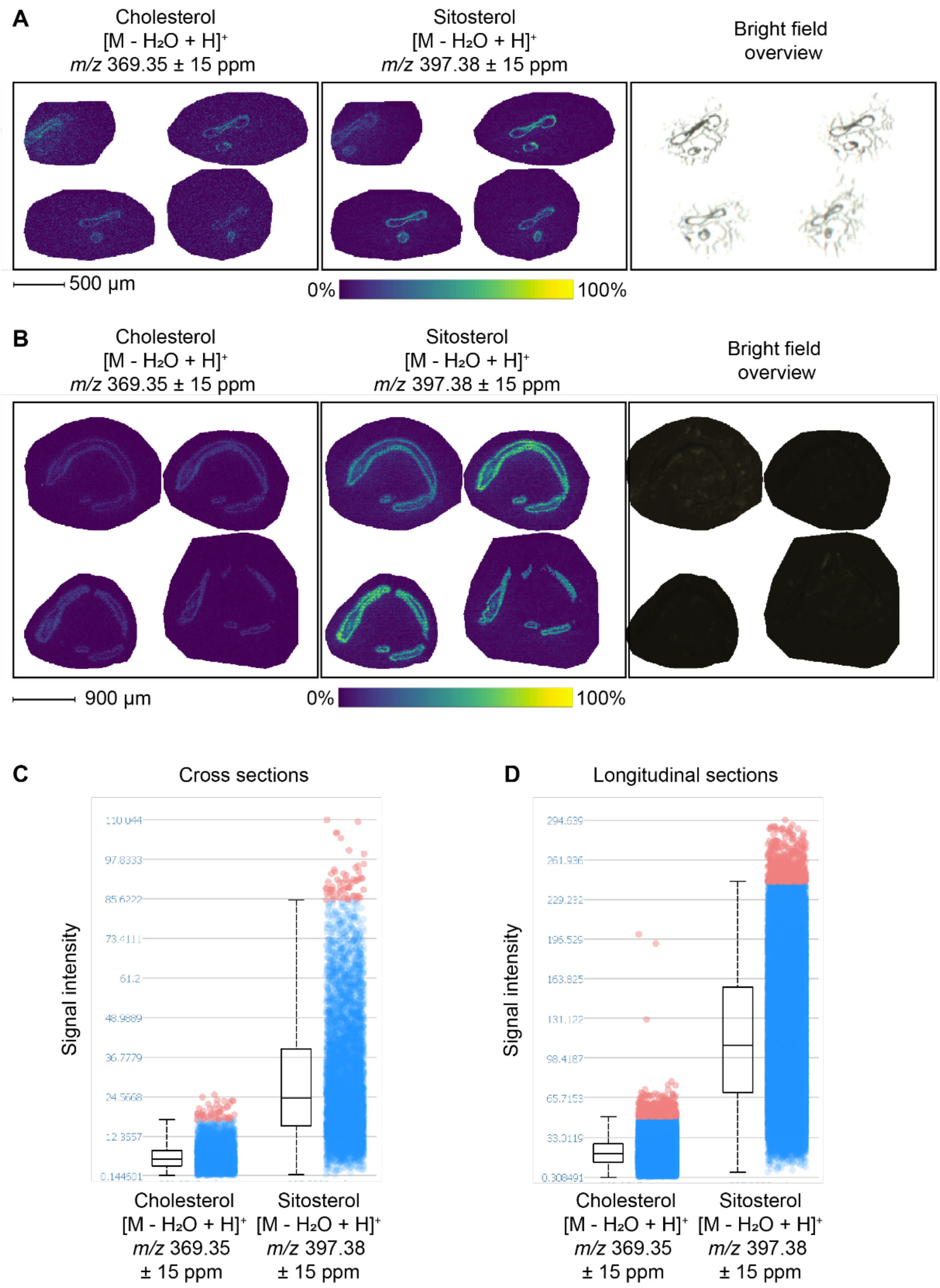

**Supplementary Figure 1 | Sitosterol and cholesterol are evenly distributed throughout *Olavius algarvensis* tissues.** Panels show cholesterol ([M-H2O+H]$^+$ at *m/z* 369.352) and sitosterol ([M-H2O+H]$^+$ at *m/z* 397.382) distribution in *O. algarvensis* cross sections (**A**) and longitudinal sections (**B**) measured by MALDI-2-MSI. Sitosterol is more abundant than cholesterol in the worm tissues (**C** and **D**).

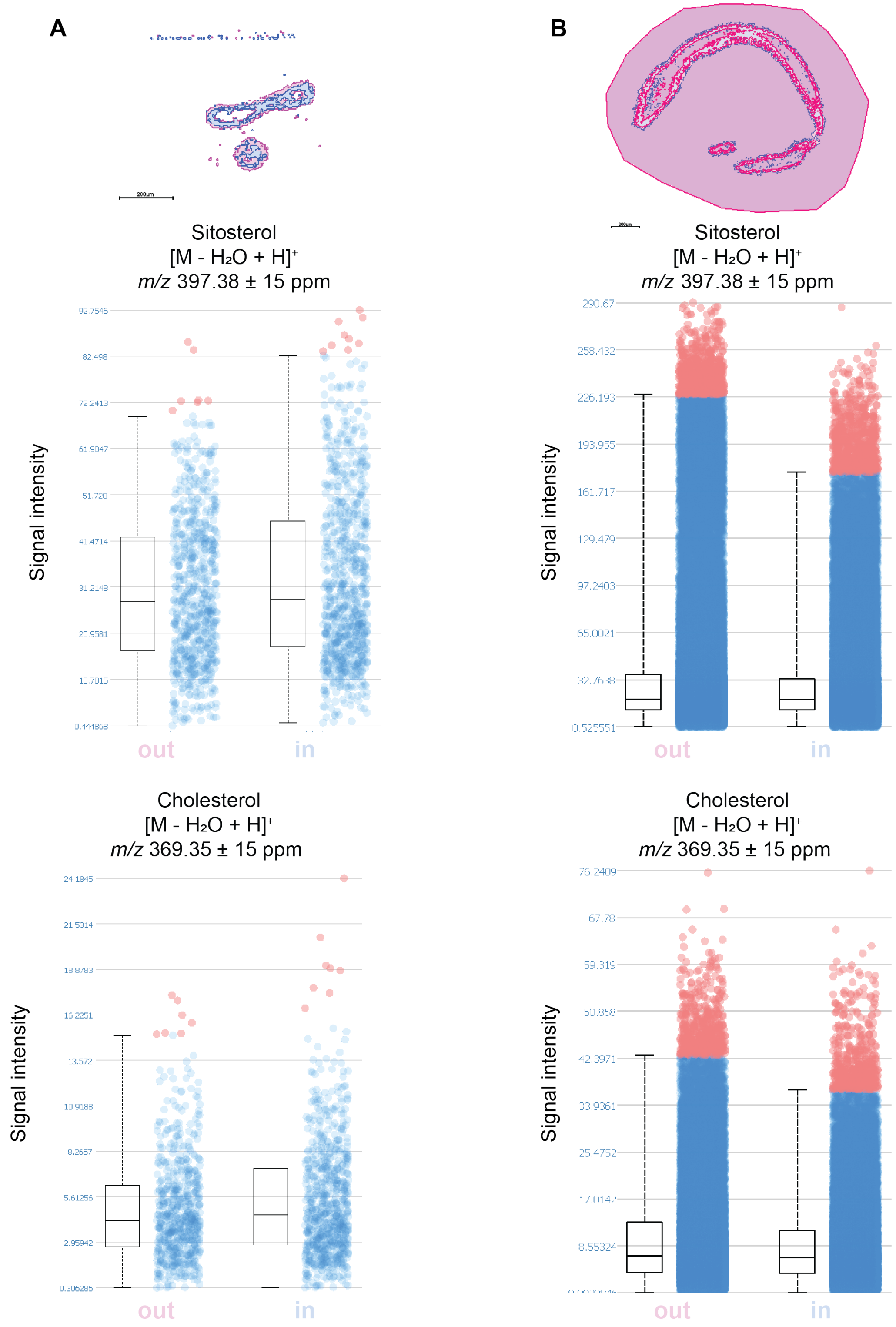

**Supplementary Figure 2 | No tissue-specific distribution of cholesterol and sitosterol in *Olavius algarvensis*.** Both sterols were uniformly distributed throughout the animals' tissues. Distribution of cholesterol and sitosterol as measured with MALDI-2-MSI. The data was segmented and the signal intensity of both sterols measured to investigate tissue specific sterol distribution in cross sections (**A**) and longitudinal sections (**B**). The colonized tissues (out) do not contain more sterols than the host-only tissues (in). The presence of the symbionts does not influence the sterol distribution in the host tissues.

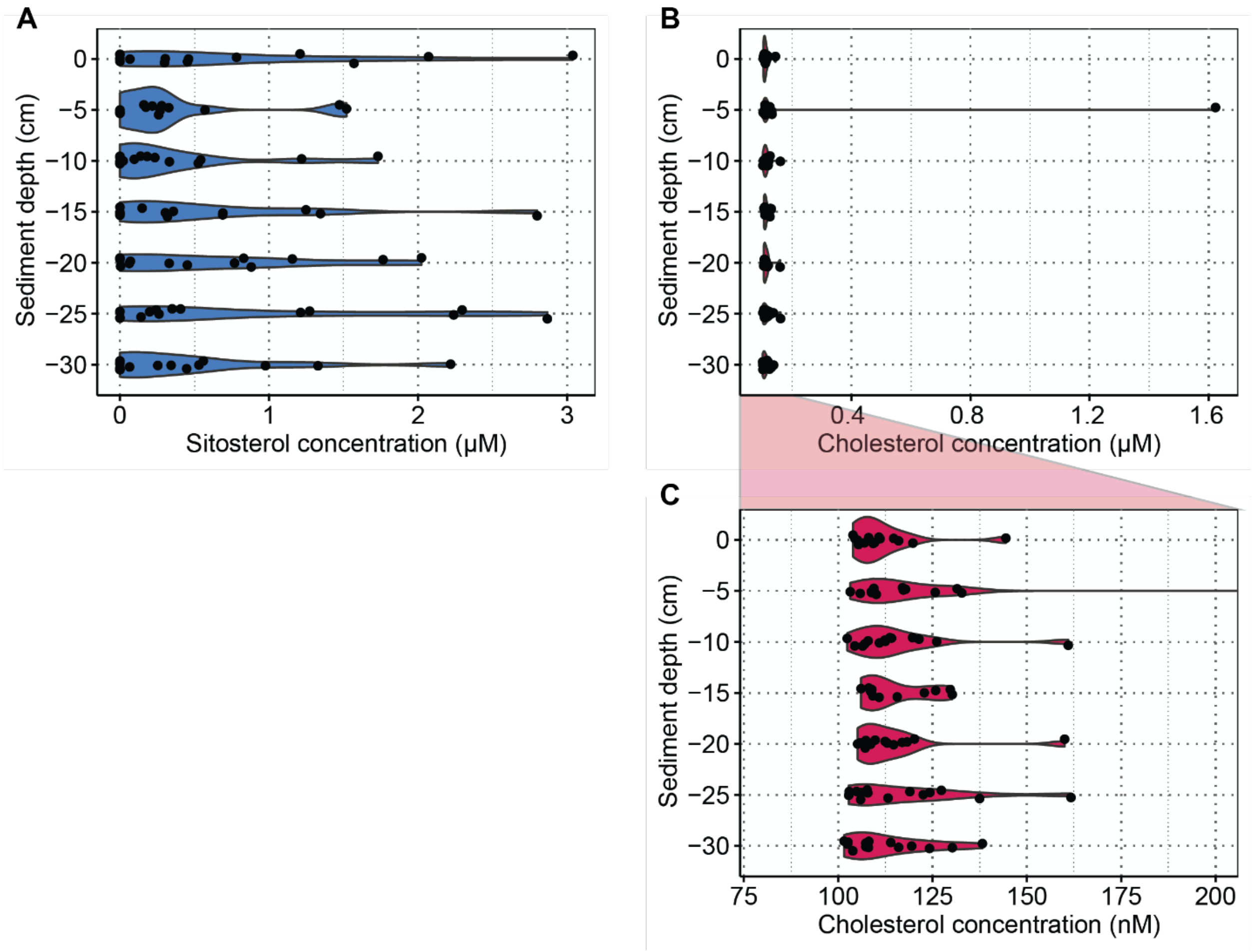

**Supplementary Figure 3 | Sediment porewater below a *Posidonia oceanica* meadow (Elba, Italy) contained sitosterol and cholesterol. A**, Sitosterol (blue) was present at micromolar concentration under the seagrass bed. **B,** Cholesterol (red) was present in nanomolar concentrations under the seagrass bed, with one exception in the low micromolar range. **C,** Magnified view of **B** to show the variation in cholesterol concentrations between 100 to 200 nanomolar.

A)

Squalene
↓

| Enzyme | Genome | Transcriptome | Proteome | Function | E.C. number |
|---|---|---|---|---|---|
| SQE | ● | ● | | Squalene monooxygenation | 1.14.14.17 |
| LAS | ● | ● | ● | Oxydosqualene cyclization | 5.4.99.7 and 5.4.99.8 |
| CYP51 | ● | ● | ● | C-14 demethylation | 1.14.14.154 |
| LBR | ● | ● | ● | C-14 reduction | 1.3.1.70 |
| MSMO1 | ● | ● | | C-4 demethylation | 1.14.18.9 /10 /11 |
| NSDHL | ● | ● | ● | | 1.1.1.170 and 1.1.1.418 |
| HSD17B7 | ● | ● | | | 1.1.1.270 |
| EBP | ● | ● | ● | Δ7-Δ8 isomerization | 5.-.-.- and 5.3.3.5 |
| SC5DL | ● | ● | | C-5 desaturation | 1.14.19.20 and 1.14.21.6 |
| DHCR7 | ● | ● | | Δ7 reduction | 1.3.1.21 |
| DHCR24 | ● | ● | | Δ24 reduction | 1.3.1.71/72 and 1.3.1.M20 |

↓
Cholesterol

B)

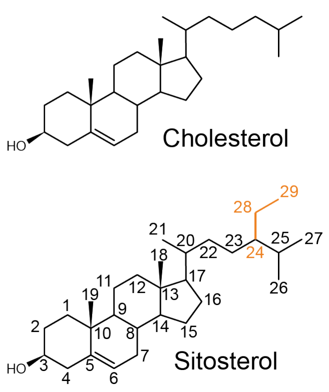

**Supplementary Figure 4 | *Olavius algarvensis* encodes and expresses all the enzymes needed to synthesize sitosterol from an intermediate of the cholesterol synthesis pathway**. **A,** All 11 enzymes involved in cholesterol synthesis were encoded and expressed by *O. algarvensis*. Dots indicate detection in the genome, transcriptome and proteome of *O. algarvensis*. **B,** An ethyl group attached to the sterol side chain differentiates the phytosterol sitosterol from the animal sterol cholesterol.

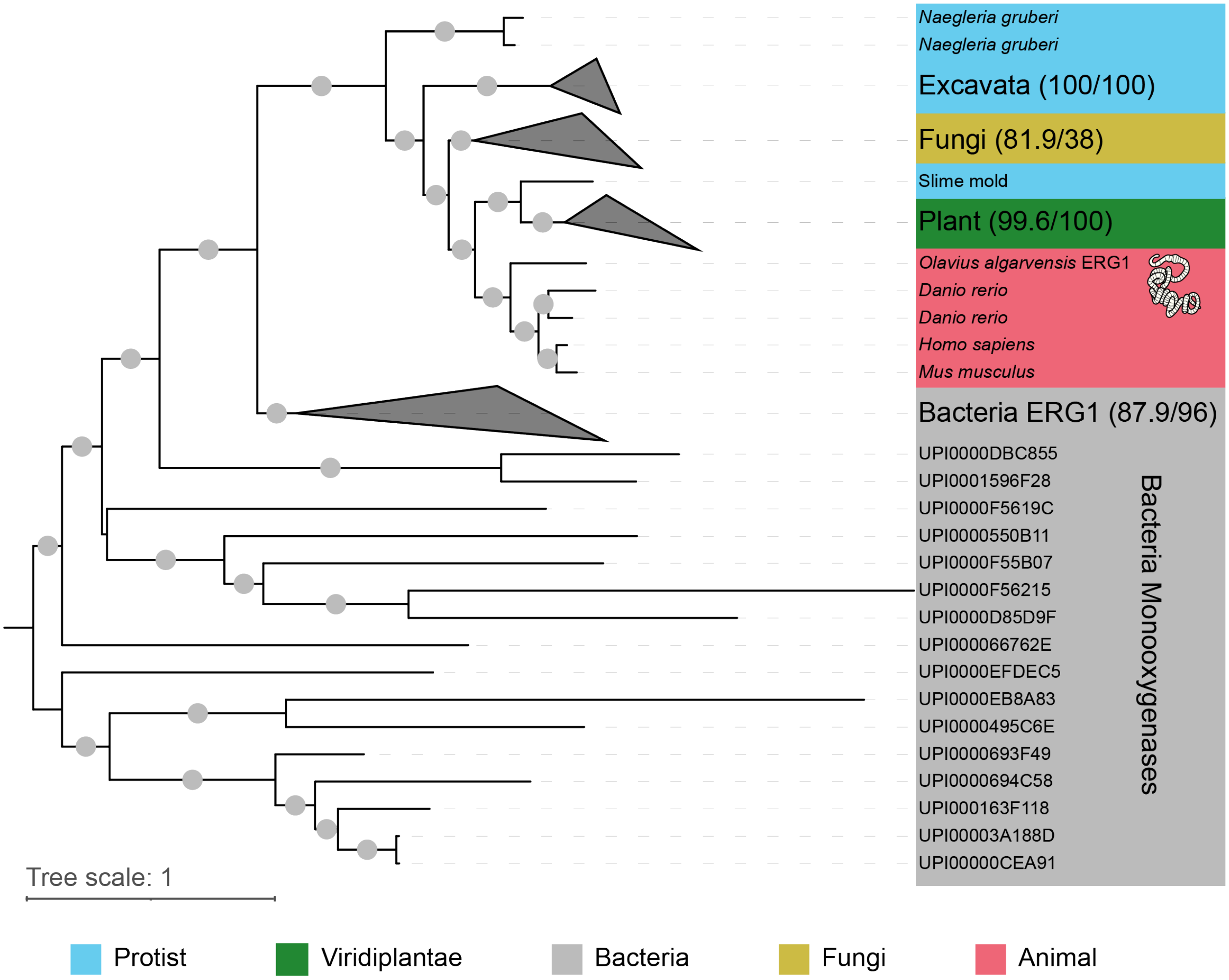

**Supplementary Figure 5 | *Olavius algarvensis* squalene monooxygenase (ERG1) sequence clusters with animal sequences.** Maximum likelihood tree of ERG1 amino acid sequences. Bootstrap values ≥90% are shown with a grey circle. The bootstrap value and branch support value are indicated in brackets for collapsed groups. The tree was rooted at midpoint in iTOL.

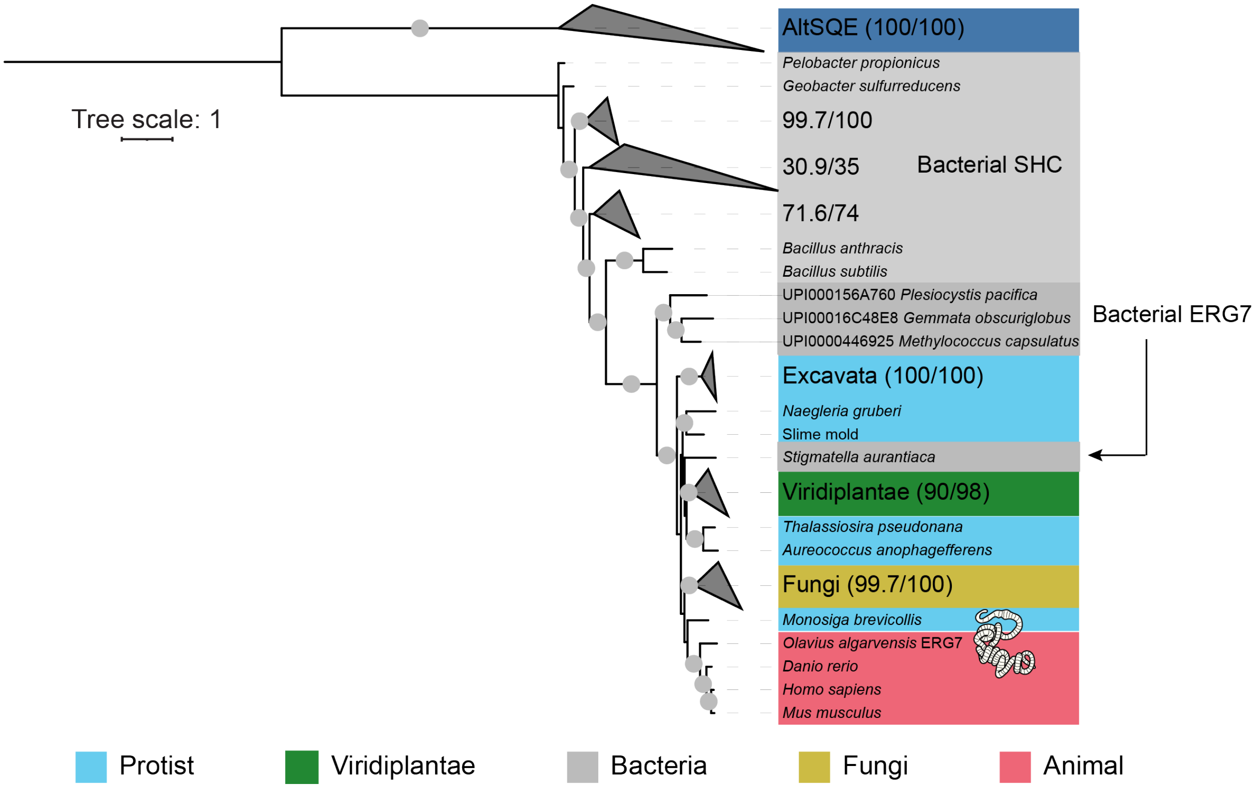

**Supplementary Figure 6 | *Olavius algarvensis* lanosterol synthase (ERG7) sequence clusters with animal sequences.** Maximum likelihood tree of ERG7 amino acid sequences, alternative squalene epoxidase (AltSQE) and the bacterial squalene-hopene cyclase (SHC) were used as outgroups. Bootstrap values ≥90% are shown with a grey circle. The bootstrap value and branch support value are indicated in brackets for collapsed groups. The tree was rooted at midpoint in iTOL.

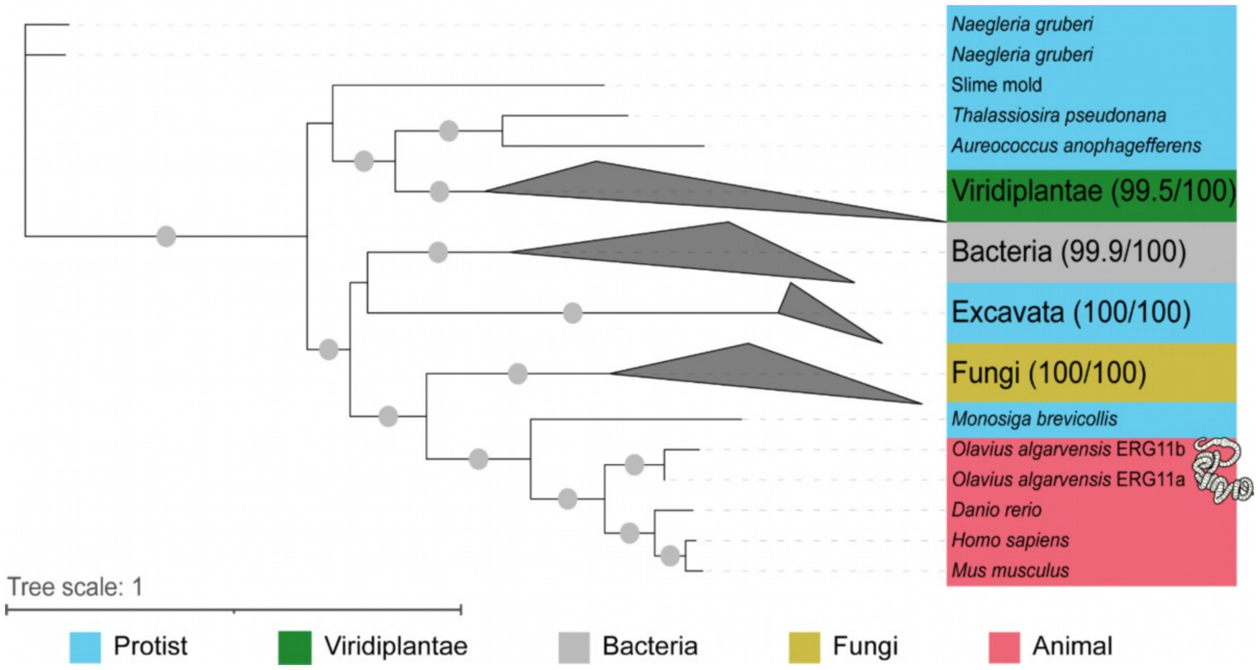

**Supplementary Figure 7 | *Olavius algarvensis* sterol delta-14-demethylase (ERG11) sequences cluster with animal sequences.** Maximum likelihood tree of ERG11 amino acid sequences. Bootstrap values ≥90% are marked with a grey circle. The bootstrap value and branch support value are indicated in brackets for collapsed groups. The tree was rooted at midpoint in iTOL.

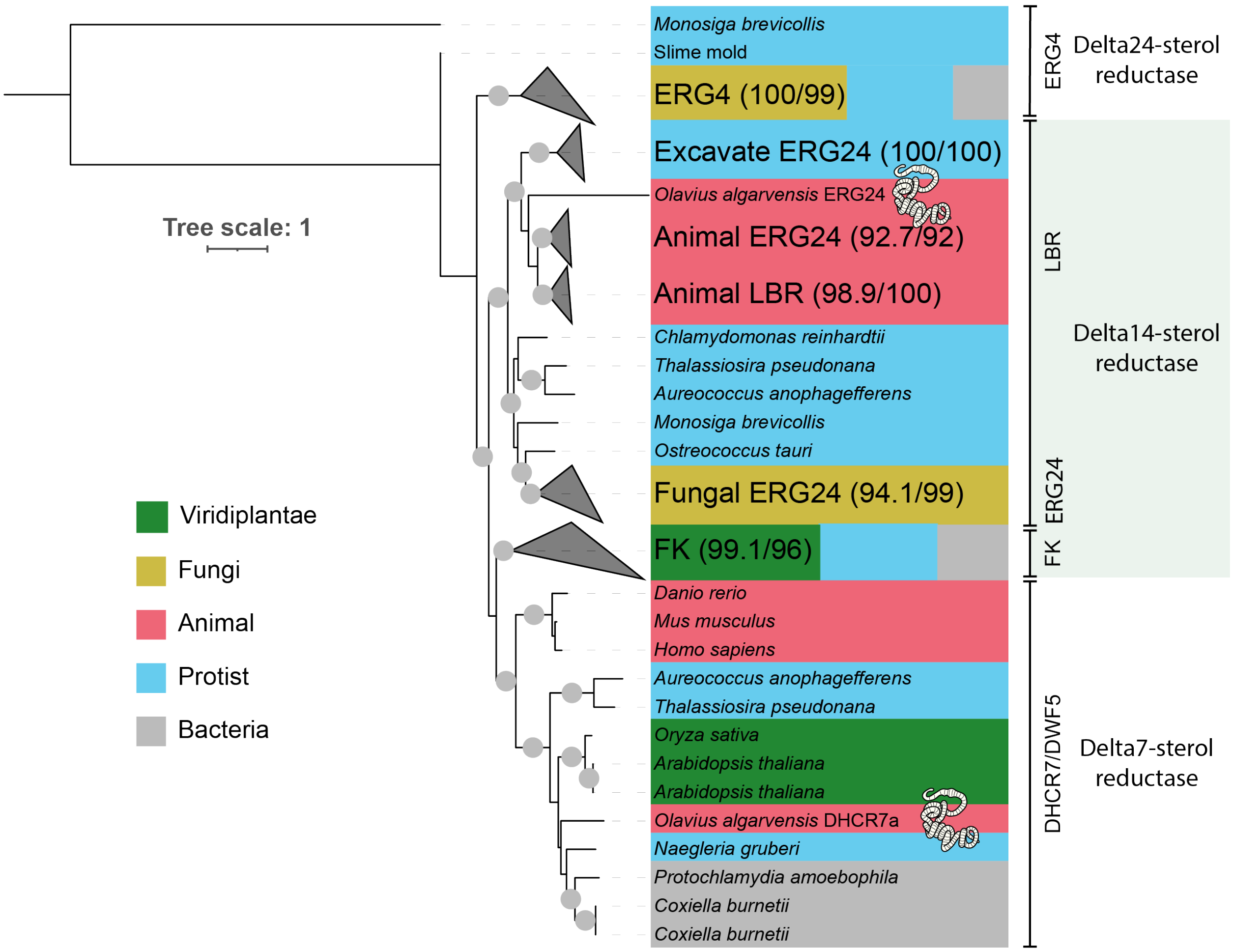

**Supplementary Figure 8 | *Olavius algarvensis* delta14-sterol reductase (ERG24) sequence clusters with animal sequences while the sterol 7-dehydrocholesterol reductase (DHCR7) cluster with plants and unicellular sequences.** The placement of *O. algarvensis* DHCR7 might indicate affinity for a substrate different from that of other animals. Maximum likelihood tree of the (ERG4)-(DHCR7/DWF5)-(ERG24/LBR/FK) protein family, here we have an example of enzyme performing different steps in fungi, animals and plant but that are evolutionary related. Bootstrap values ≥90% are marked with a grey circle. The bootstrap value and branch support value are indicated in brackets for collapsed groups. The tree was rooted at midpoint in iTOL.

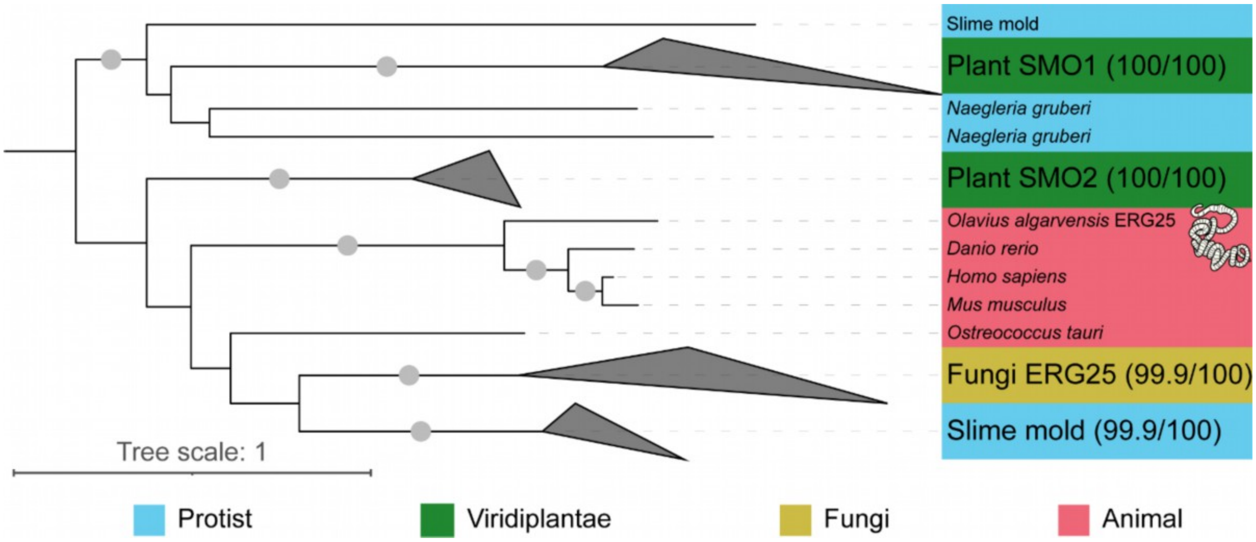

**Supplementary Figure 9 | *Olavius algarvensis* methylsterol monooxygenase sequence clusters with animal sequences.** Maximum likelihood tree of methylsterol monooxygenase (SMO1/SMO2-ERG25-MSMO) amino acid sequences. Bootstrap values ≥90% are marked with a grey circle. The bootstrap value and branch support value are indicated in brackets for collapsed groups. The tree was rooted at midpoint in iTOL.

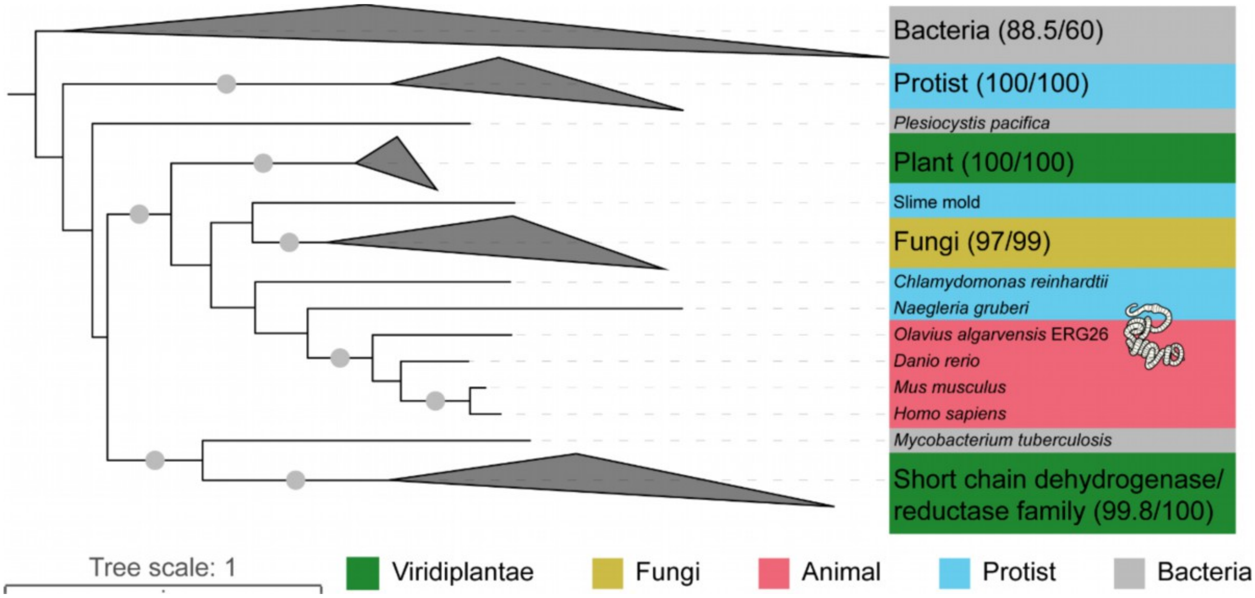

**Supplementary Figure 10 | *Olavius algarvensis* sterol-4-alpha-carboxylate 3-dehydrogenase, decarboxylating sequence clusters with animal sequences.** Maximum likelihood tree of sterol-4-alpha-carboxylate 3-dehydrogenase (HSD-NSDHL-ERG26) amino acid sequences. Bootstrap values ≥90% are marked with a grey circle. The bootstrap value and branch support value are indicated in brackets for collapsed groups. The tree was rooted at midpoint in iTOL.

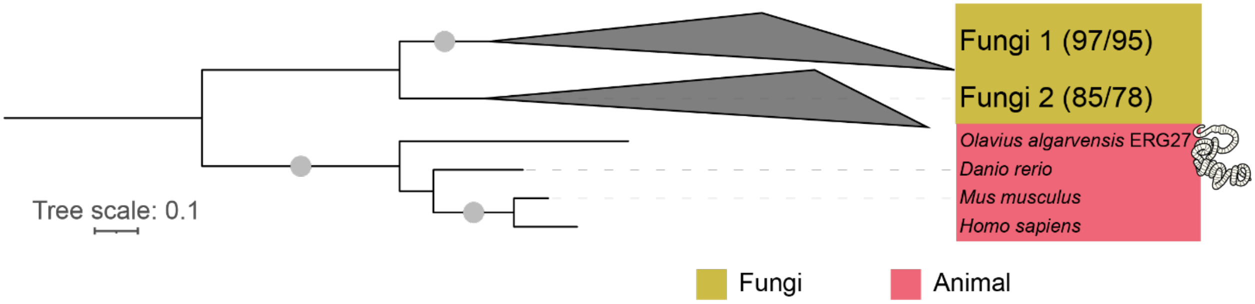

**Supplementary Figure 11 | *Olavius algarvensis* 3-keto reductase (ERG27) sequence clusters with animal sequences.** Maximum likelihood tree of ERG27 amino acid sequences. Bootstrap values ≥90% are marked with a grey circle. The bootstrap value and branch support value are indicated in brackets for collapsed groups. The tree was rooted at midpoint in iTOL. The gene performing C-3 keto reduction in plants and protists is still unknown.

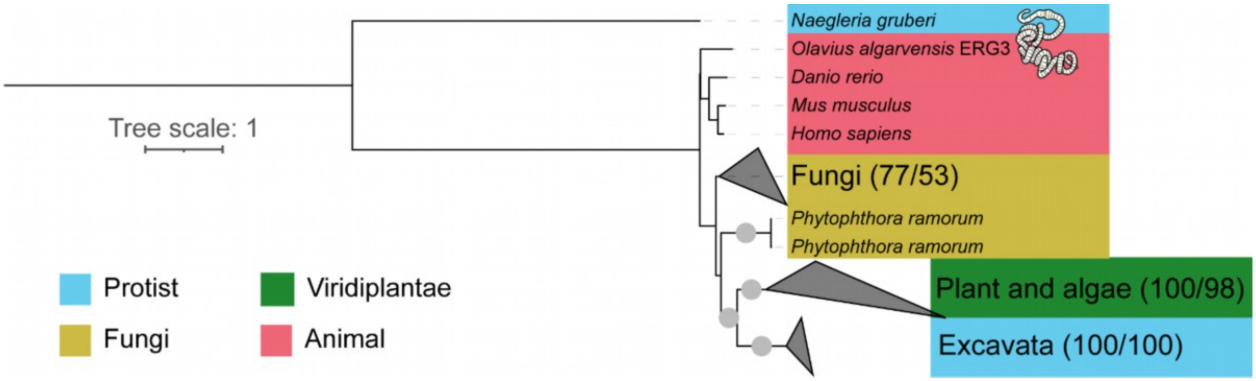

**Supplementary Figure 12 | *Olavius algarvensis* sterol C-5 desaturase (ERG3) sequence clusters with animal sequences.** Maximum likelihood tree of ERG3 amino acid sequences. Bootstrap values ≥90% are marked with a grey circle. The bootstrap value and branch support value are indicated in brackets for collapsed groups. The tree was rooted at midpoint in iTOL.

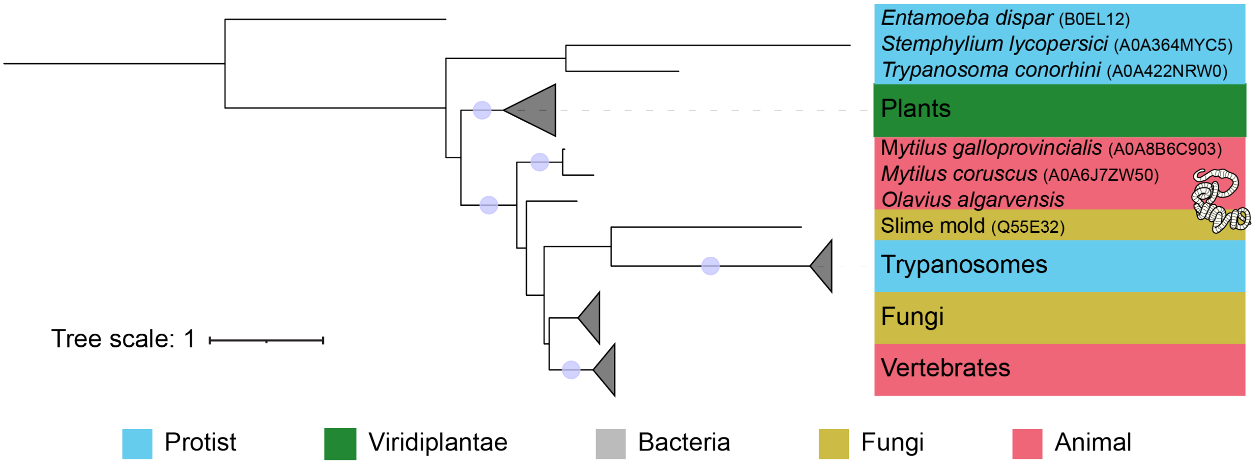

**Supplementary Figure 13 | *Olavius algarvensis* detla7-delta8-sterol isomerase (EBP) sequence clusters with animal and fungal sequences.** Maximum likelihood tree of EBP amino acid sequences. Bootstrap values ≥95% are marked with a grey circle. The tree was rooted at midpoint in iTOL.

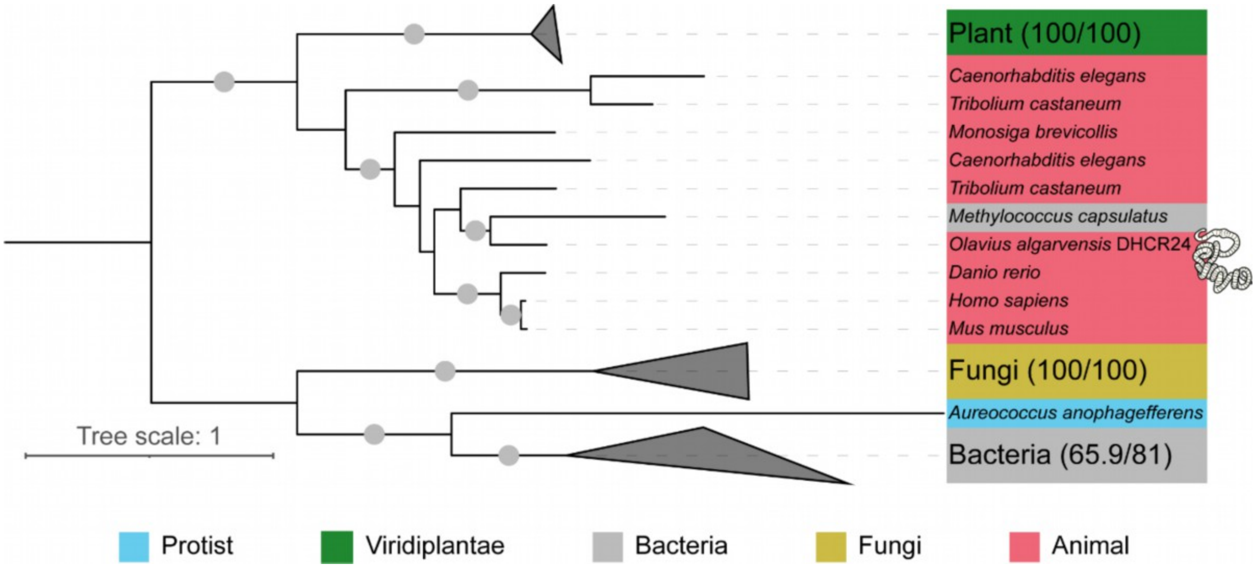

**Supplementary Figure 14 | *Olavius algarvensis* sterol 24-dehydrocholesterol reductase (DHCR24) sequence clusters with animal sequences.** Maximum likelihood tree of DHCR24 amino acid sequences. Bootstrap values ≥90% are marked with a grey circle. The bootstrap value and branch support value are indicated in brackets for collapsed groups. The tree was rooted at midpoint in iTOL.

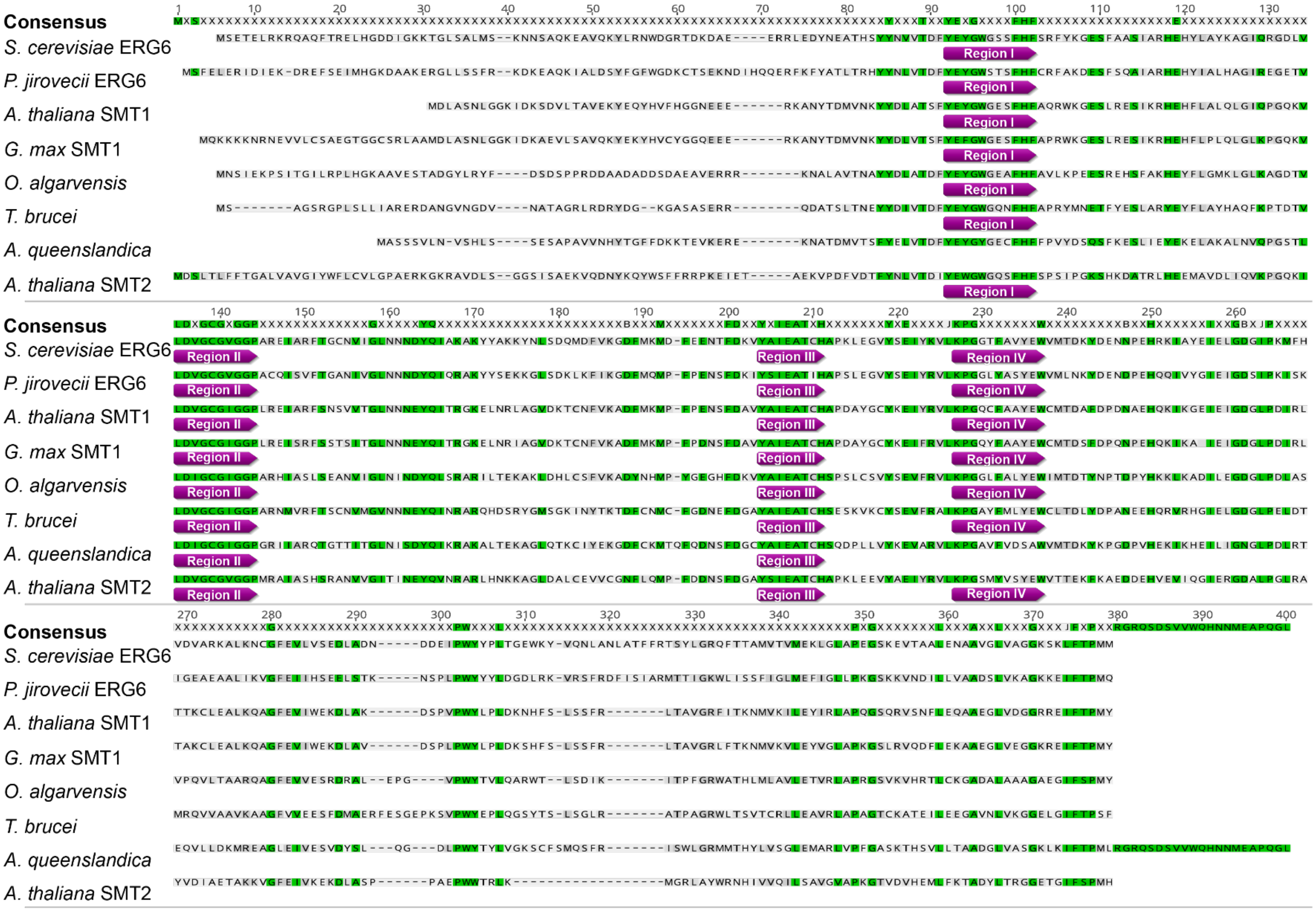

**Supplementary Figure 15 | Alignment of C-24 sterol methyltransferase ($C_{24}$-SMT) amino acid sequences.** Sequences from: fungi, *Saccharomyces cerevisiae* (P25087) and *Pneumocystis* (Q96WX4); plants: *Arabidopsis thaliana* SMT1 (Q9LM02) and SMT2 (Q39227) and *Glycine max* SMT1 (Q43445); gutless annelid: *Olavius algarvensis* (this study); excavate: *Trypanosoma brucei* (Q4FKJ2); and sponge: *Amphimedon queenslandica* (A0A1X7ULF8). The sequences were aligned using ClustalW (Geneious). Purple labels indicate sterol (II) and AdoMet (I, III and IV) binding regions.

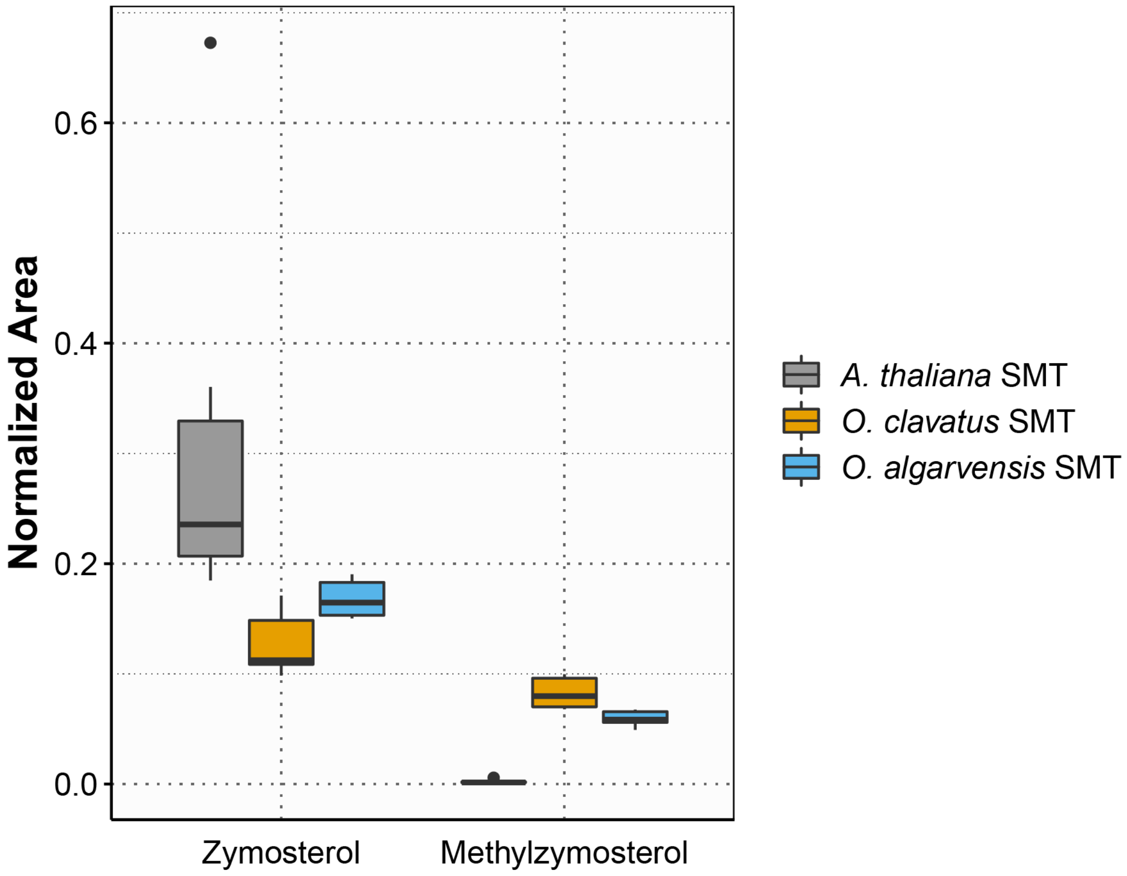

**Supplementary Figure 16 | The $C_{24}$-SMT of *Olavius spp.* used zymosterol as a substrate for the first methylation.** . *Arabidopsis thaliana* $C_{24}$-SMT did not methylate zymosterol. The heterologously expressed $C_{24}$-SMT enzymes from both *O. algarvensis* and *O. clavatus* were able to methylate zymosterol to methylzymosterol. The peak of the substrate (zymosterol) and of the product (methylzymosterol) were integrated at the end of the enzymatic assay, in which *E. coli* expressed $C_{24}$-SMT from each taxon ($n$ = 5 in each case) was incubated with zymosterol as substrate. The abundance was normalized with an internal standard.

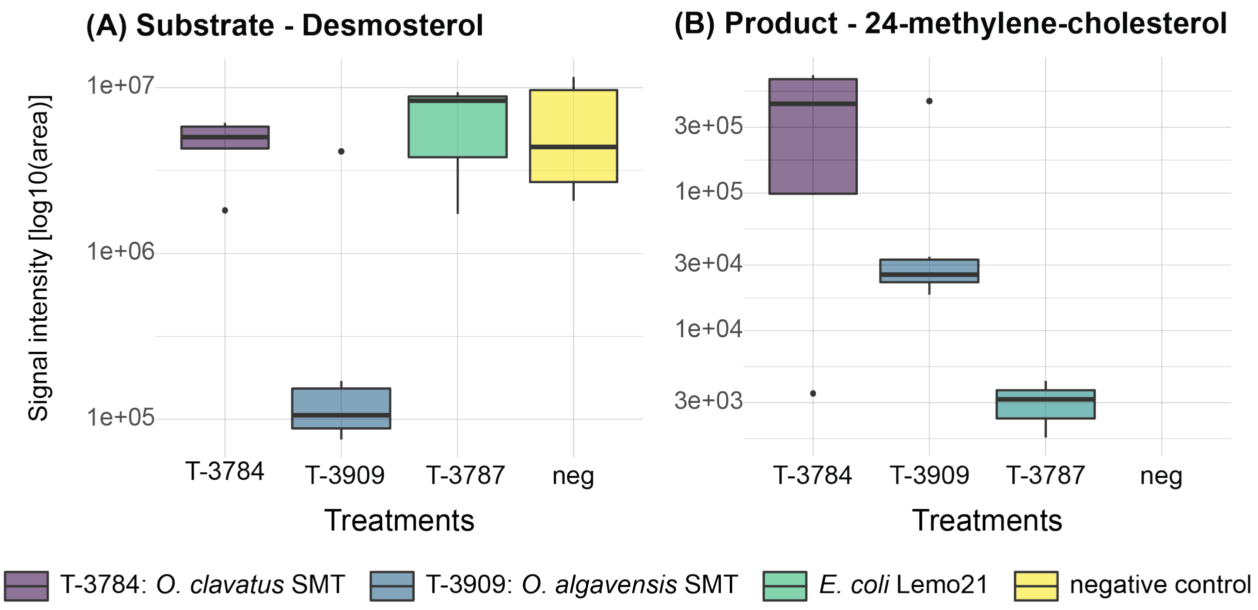

**Supplementary Figure 17 | The $C_{24}$-SMT of *Olavius spp*. used desmosterol as a substrate for the first methylation step.** *Arabidopsis thaliana* $C_{24}$-SMT was not able to methylate desmosterol. The heterologously expressed $C_{24}$-SMT enzymes from both *O. algarvensis* and *O. clavatus* were able to methylate desmosterol to produce 24-methylene-cholesterol. The abundance of **(A)** the substrate (desmosterol) and **(B)** the product (24-methylene-cholesterol) were measured at the end of the enzymatic assay, in which *E. coli* expressed $C_{24}$-SMT from each taxon ($n$ = 5 in each case) was incubated with desmosterol as substrate.

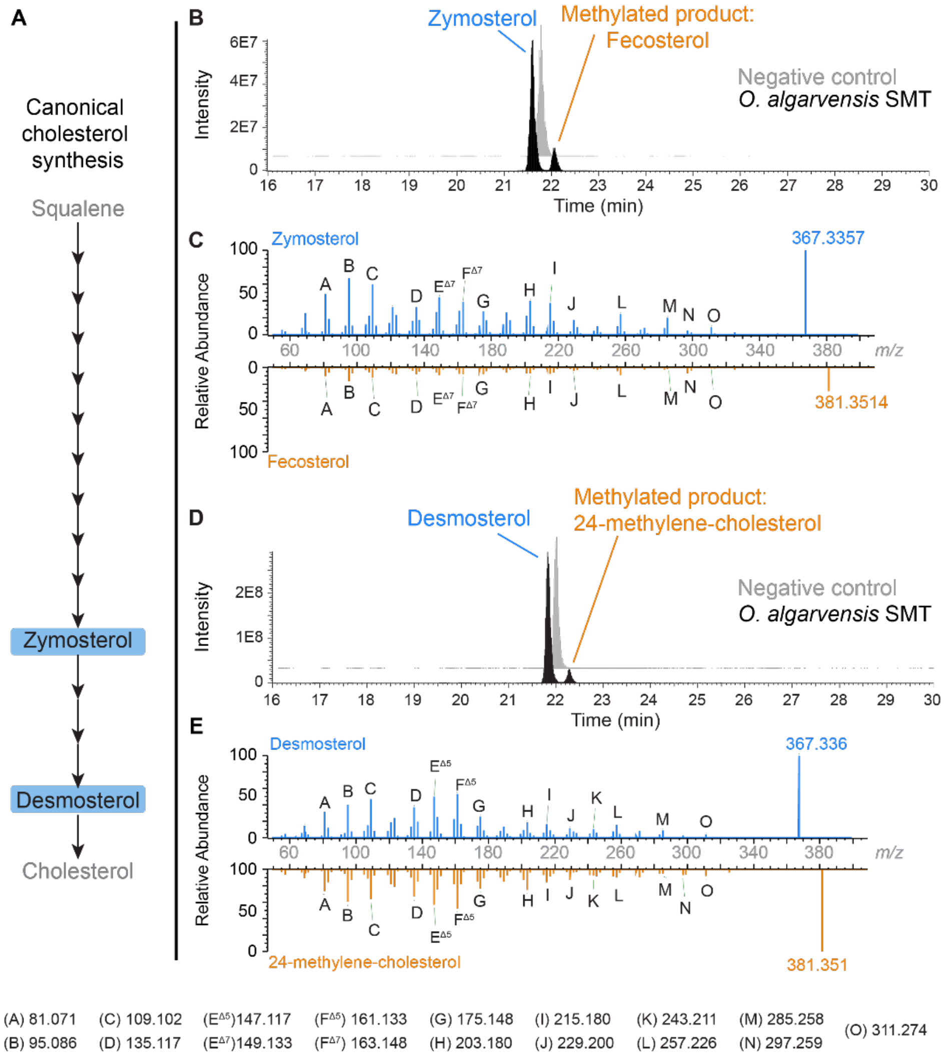

**Supplementary Figure 18 | $C_{24}$-SMT of *Olavius algarvensis* uses two intermediates of the cholesterol synthesis, zymosterol and desmosterol, as substrates for methylation. A,** Zymosterol and desmosterol are intermediates of the classical animal cholesterol synthesis. They are produced in the second half of the cholesterol synthesis pathway. **B and D,** *O. algarvensis* $C_{24}$-SMT, after overexpression in *E. coli*, added a methyl group to the side chain of zymosterol and desmosterol. LC-MS chromatograms of the enzymatic assay performed with zymosterol and desmosterol as substrates**. C and E,** The substrates and methylated products were identified by MS/MS.

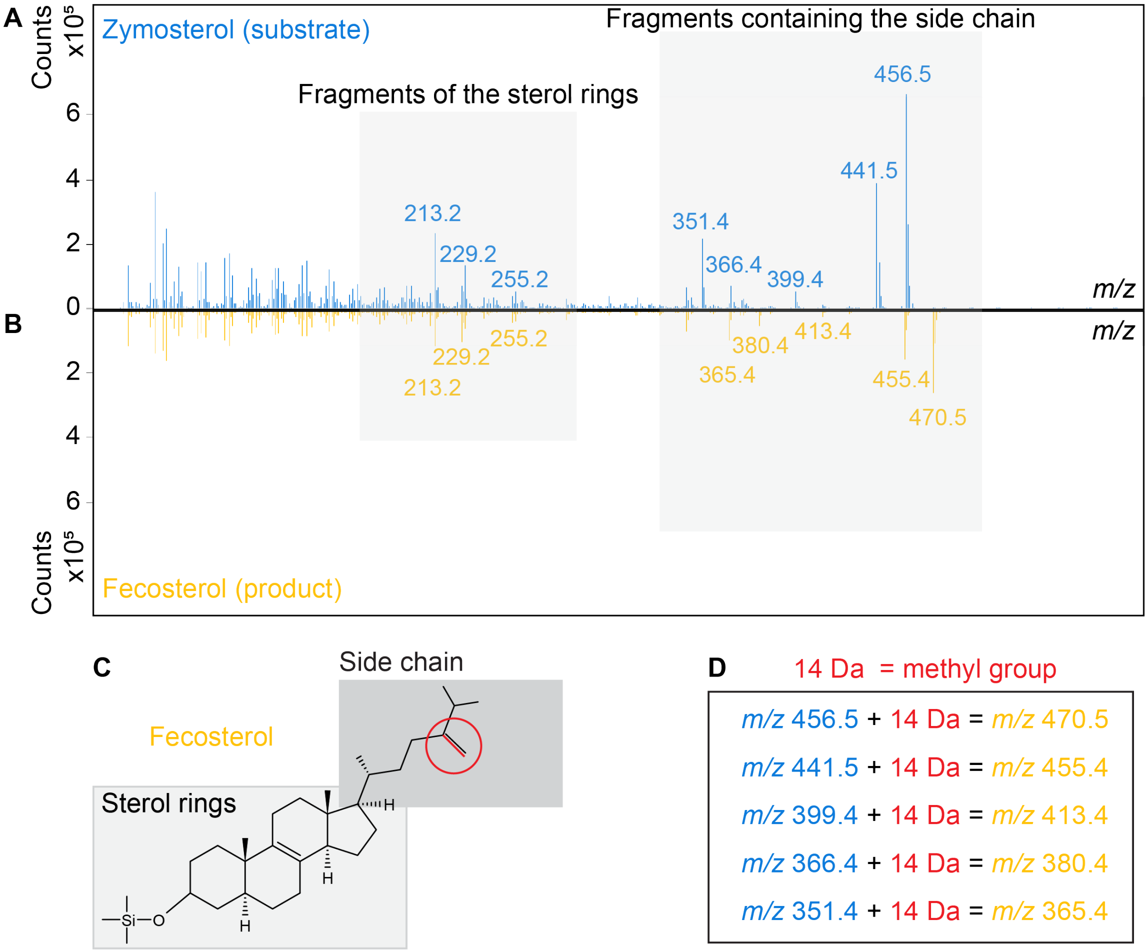

**Supplementary Figure 19 | The $C_{24}$-SMT of *Olavius algarvensis* added a methyl group to the side chain of zymosterol. A, B,** Representative mass spectra of zymosterol **(A)** and the compound identified as fecosterol (24-methylene-zymosterol) **(B). C,** Structure of fecosterol, with the methyl group highlighted in red. **D,** In the comparison of the mass spectra of zymosterol and fecosterol, the fragments containing the side chain were shifted by 14 Da, a difference that represents the addition of a methyl group. No mass shift was observed in the sterol ring fragments. These results indicate that a methyl group was added to the zymosterol side chain.

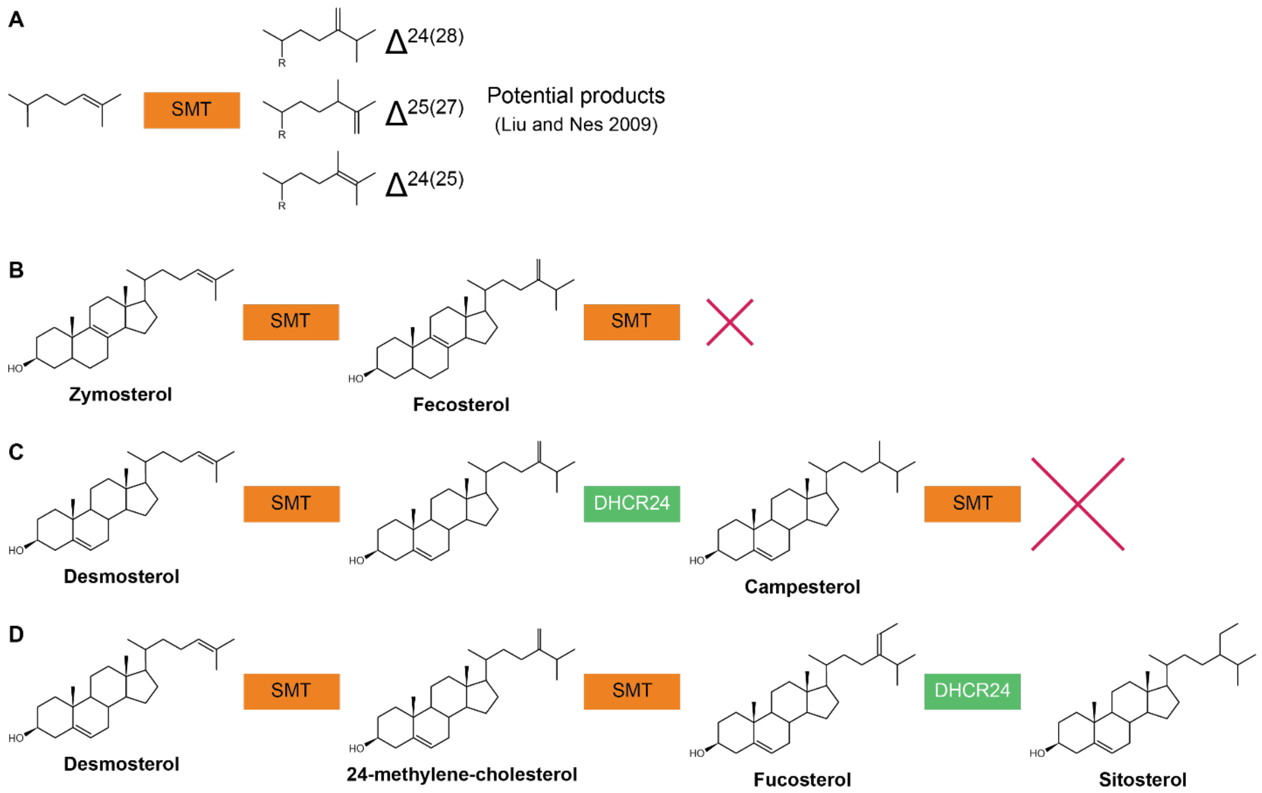

**Supplementary Figure 20 | Potential substrates for the second $C_1$-transfer based on the results of the first $C_1$-transfer. A,** $C_{24}$-SMT can produce different products. *O. algarvensis* $C_{24}$-SMT mainly produced methylene ($\Delta^{24(28)}$) products. **B,** *O. algarvensis* $C_{24}$-SMT methylates zymosterol to produce fecosterol. Fecosterol, product of the first methylation, was not methylated further by *O. algarvensis* $C_{24}$-SMT even when the incubation time was extended. **C and D,** *O. algarvensis* $C_{24}$-SMT added a methyl group to demosterol to produce 24-methylene-cholesterol. **C,** The methylene group is then likely reduced by sterol C24-reductase (DHCR24), producing campesterol. Campesterol was identified as a potential candidate for the second methylation step but *O. algarvensis* $C_{24}$-SMT could not use it as a substrate. **D,** The product of desmosterol methylation could also be used directly as a substrate for the second methylation. *O. algarvensis* $C_{24}$-SMT added a methyl group to 24-methylene-cholesterol, producing fucosterol. DHCR24, which is expressed based on its presence in *O. algarvensis* transcriptomes, could then remove the $\Delta^{24(28)}$ double bond and transform fucosterol into sitosterol.

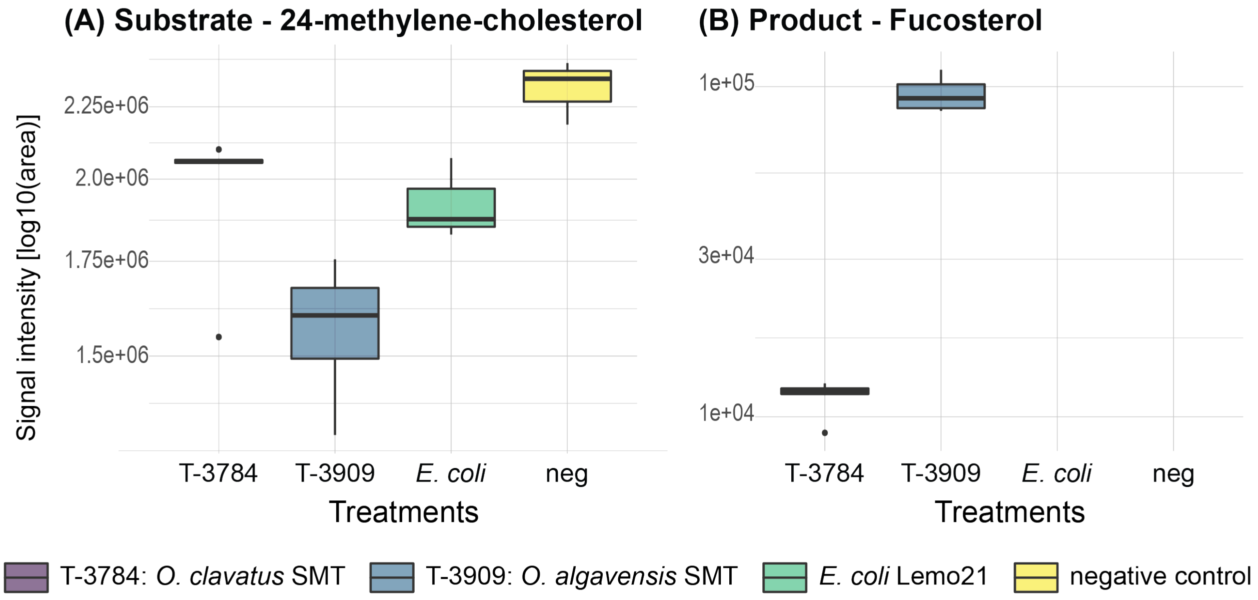

**Supplementary Figure 21 | The $C_{24}$-SMT of *Olavius* spp. use 24-methylene-cholesterol as a substrate for the second methylation reaction.** The enzymes from both *O. algarvensis* and *O. clavatus* used 24-methylene-cholesterol as substrate and produced fucosterol. The abundance of **(A)** the substrate (24-methylene-cholesterol) and **(B)** the product (fucosterol) were integrated at the endpoint of the enzymatic assay, in which *E. coli* expressed $C_{24}$-SMT from each taxon ($n$ = 5 in each case) was incubated with 24-methylene-cholesterol as substrate.

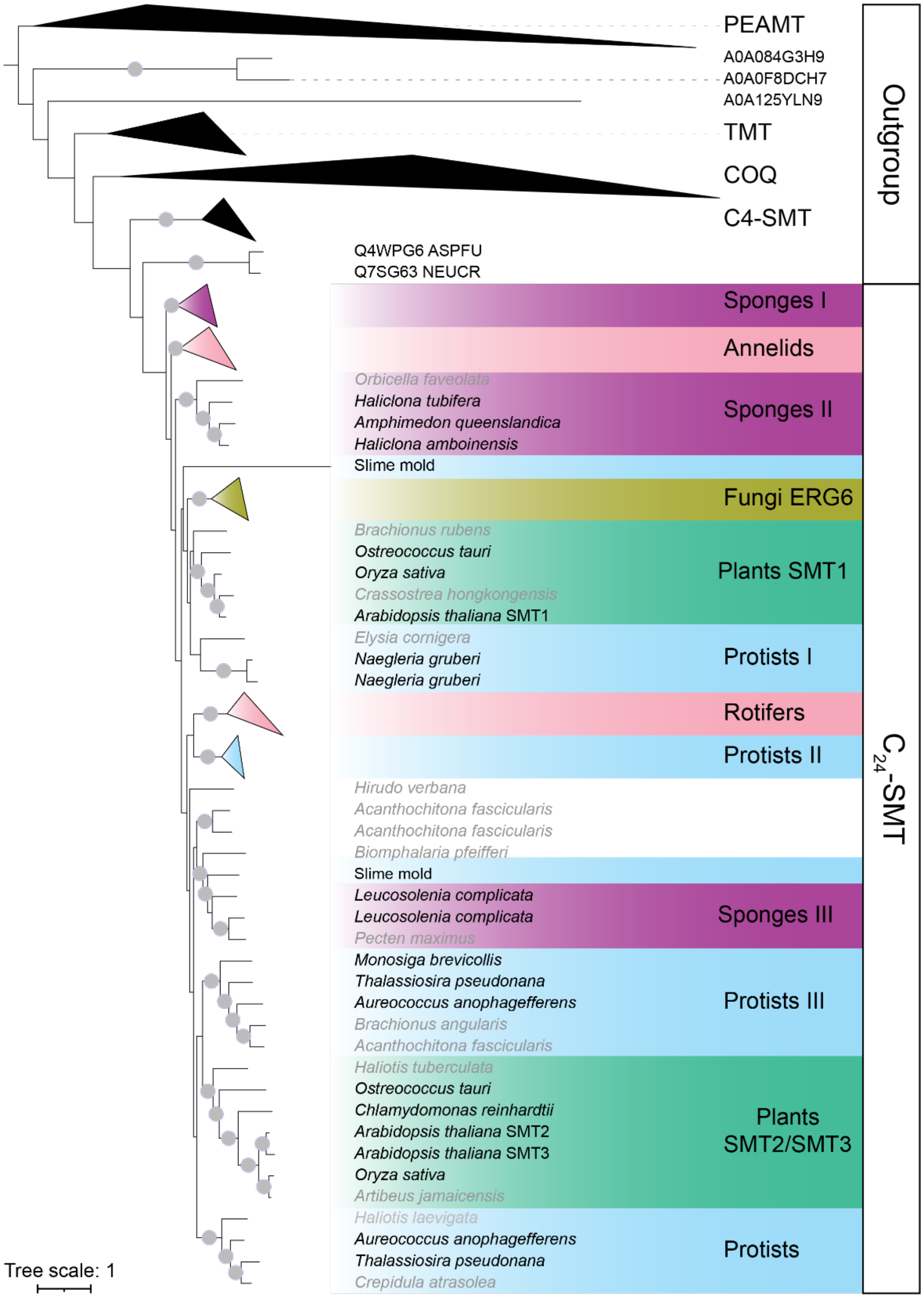

**Supplementary Figure 22 | Most C-24 sterol methyltransferase ($C_{24}$-SMT) homologues identified by BLAST within mollusks, chordata and nematode phyla were plant, protist or fungal contaminations, or belonged to the $C_4$-SMT, an SMT specific to nematodes.** Maximum likelihood amino acid tree of SMTs, with other SAM-dependent methyltransferases used as outgroups. Bootstrap values ≥95% are marked with a grey circle. Sequences isolated from animal samples but identified as contamination are labelled in grey. The tree was rooted at midpoint in iTOL. Ubiquinone biosynthesis O-methyltransferase (COQ), phosphoethanolamine N-methyltransferase (PEAMT), tocopherol O-methyltransferase (TMT), C-4 sterol methyltransferase ($C_4$-SMT).

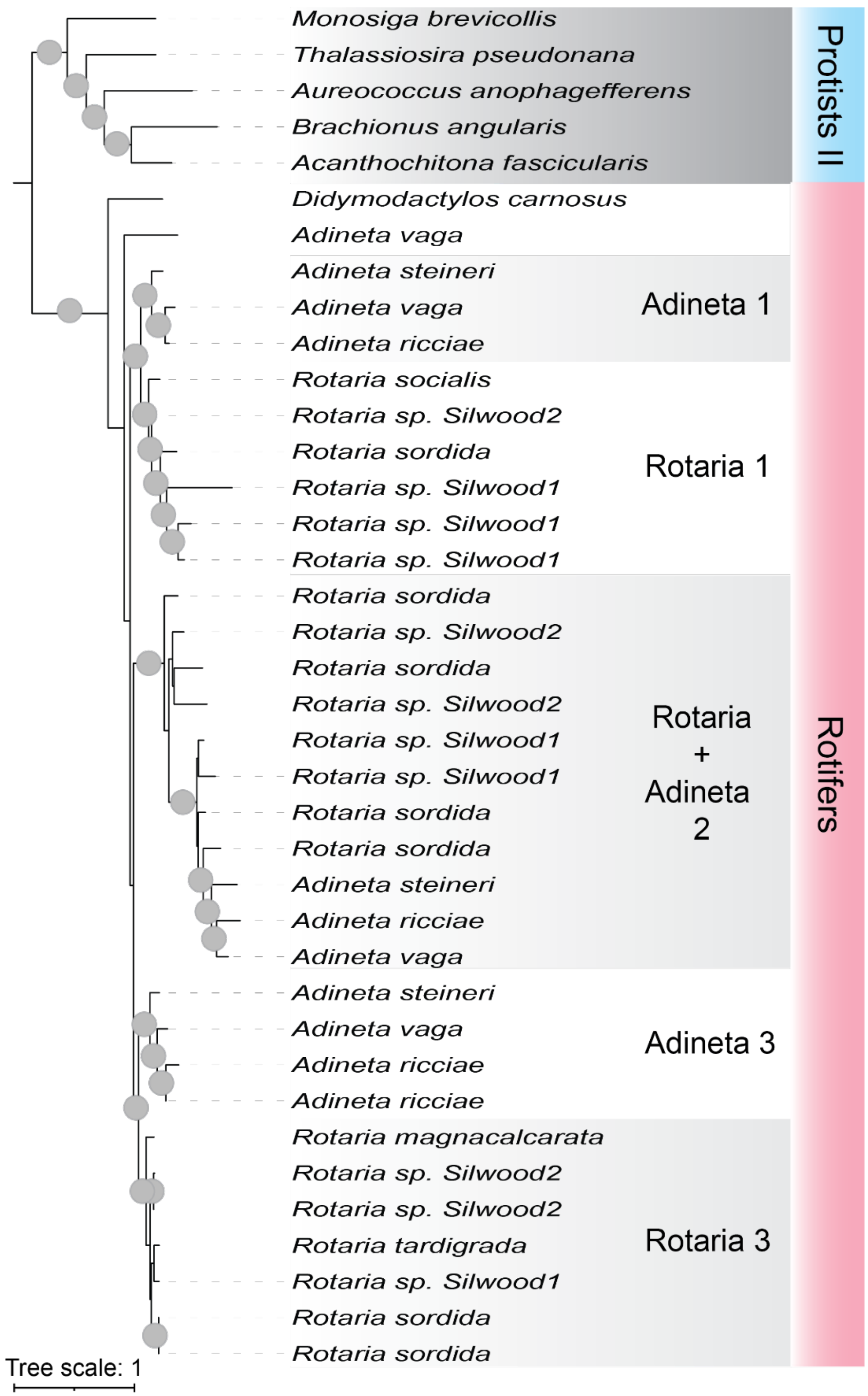

**Supplementary Figure 23 | Rotifers express several isoforms of C-24 sterol methyltransferase ($C_{24}$-SMT).** C24-SMT homologs were detected in nine species of bdeloid rotifers. Rotifers can express up to three different $C_{24}$-SMT isoforms. Zoom in on the protists II and rotifers group presented in **Supplementary Figure 22**.

### Supplementary Tables

Supplementary Table 15 | Solvent gradient for high-resolution LC-MS/MS with C18 column. .. 37

**Supplementary Table 1 | List of sterols detected by MALDI-2-MSI in *Olavius algarvensis*.**

| **Sterol** | **Sum formula** | **Molecular weight M** (calc) | **$[M-H_2O+H]^+$** (calc) | **$[M-H_2O+H]^+$** (exp) | **Mass error** (ppm) |
|---|---|---|---|---|---|
| cholesterol | $C_{27}H_{46}O$ | 386.355 | 369.352 | 369.345 | 19 |
| sitosterol | $C_{29}H_{50}O$ | 414.386 | 397.383 | 397.376 | 18 |
| stigmasterol | $C_{29}H_{48}O$ | 412.371 | 395.368 | 395.364 | 10 |

**Supplementary Table 2 | Detection of enzymes involved in sterol biosynthesis in the draft genome of *Olavius algarvensis*.**

| Enzyme | Target | Contig | Status | Reasons | Matches | Mismatch | ID | Coverage | Score | # of exons |
|---|---|---|---|---|---|---|---|---|---|---|
| SQE | 4410_host_Oalg_flye-2.8 | contig_6471 | incomplete | missing stopcodon + missmatch | 515 | 3 | 99.4 | 100 | 0.988 | 9 |
| LAS | 4410_host_Oalg_flye-2.8 | contig_13849 | auto | - | 756 | 0 | 100 | 100 | 1 | 20 |
| CYP51 | 4410_host_Oalg_flye-2.8 | contig_37932 | incomplete | mismatches | 504 | 2 | 99.6 | 100 | 0.992 | 11 |
| CYP51 | 4410_host_Oalg_flye-2.8 | contig_37932 | incomplete | mismatches | 319 | 2 | 99.4 | 100 | 0.988 | 6 |
| LBR | 4410_host_Oalg_flye-2.8 | contig_20025 | partial 1-21 (#1/2) | - | 21 | 0 | 100 | 3.6 | 0.036 | 1 |
| LBR | 4410_host_Oalg_flye-2.8 | contig_42201 | partial 22-583 (#2/2) | - | 555 | 7 | 98.8 | 100 | 0.94 | 10 |
| MSMO1 | 4410_host_Oalg_flye-2.8 | contig_24678 | incomplete | gap to querystart (7-306) | 300 | 0 | NA | NA | 0.98 | 6 |
| NSDHL | 4410_host_Oalg_flye-2.8 | contig_27038 | auto | - | 346 | 0 | 100 | 100 | 1 | 7 |
| HSD17B7 | 4410_host_Oalg_flye-2.8 | contig_21098 | partial 1-27 (#1/2) | - | 27 | 0 | NA | NA | 0.079 | 1 |
| HSD17B7 | 4410_host_Oalg_flye-2.8 | contig_21097 | partial 18-343 (#2/2) | mismatches + missing stopcodon | 312 | 4 | 98.7 | 100 | 0.898 | 9 |
| EBP | 4410_host_Oalg_flye-2.8 | contig_10005 | auto | - | 230 | 0 | 100 | 100 | 1 | 3 |
| SC5DL | 4410_host_Oalg_flye-2.8 | contig_6195 | incomplete | missing stopcodon | 282 | 0 | 100 | 100 | 1 | 3 |
| DHCR7 | 4410_host_Oalg_flye-2.8 | Contig_8717 | incomplete | missing stopcodon + missmatch | 482 | 1 | 99.8 | 100 | 0.996 | 3 |
| DHCR24 | 4410_host_Oalg_flye-2.8 | scaffold_4022 | incomplete | missing stopcodon + missmatch | 501 | 2 | 99.6 | 100 | 0.992 | 8 |
| SMT | 4410_host_Oalg_flye-2.8 | contig_41985 | incomplete | missing stopcodon | 165 | 0 | 100 | 100 | 1 | 4 |

**Supplementary Table 3 | List of the enzymes involved in sterol biosynthetic pathway in eukaryotic model organisms**.

| Enzymatic reaction | E.C number | Animal | Plant | Fungi |
|---|---|---|---|---|
| Squalene monooxygenation | 1.14.14.17 | SQE | SQE | ERG1 |
| Oxydosqualene cyclization | 5.4.99.7 and 5.4.99.8 | LAS | CAS | ERG7 |
| C-14 demethylation | 1.14.14.154 | CYP51 | CYP51G1 | ERG11 |
| C-14 reduction | 1.3.1.70 | LBR | FK | ERG24 |
| C-4 demethylation | 1.14.18.9 /10 /11 | MSMO1 | SMO | ERG25 |
| | 1.1.1.170 and 1.1.1.418 | NSDHL | 3β-HSD | ERG26 |
| | 1.1.1.270 | HSD17B7 | ? | ERG27 |
| Δ7-Δ8 isomerization | 5.-.-.- and 5.3.3.5 | EBP | HYD1 | ERG2 |
| C-5 desaturation | 1.14.19.20 and 1.14.21.6 | SC5DL | DWF7 | ERG3 |
| Δ7 reduction | 1.3.1.21 | DHCR7 | DWF5 | n.d. |
| Δ24 reduction | 1.3.1.71/72 and 1.3.1.M20 | DHCR24 | DIM | ERG4 |
| C-22 desaturation | 1.14.19.41 | n.d. | CYP710A | ERG5 |
| C-24 and C-28 methylation | 2.1.1.41 and 2.1.1.143 | n.d. | SMT | ERG6 |
| Cyclopropylsterol isomerization | 5.5.1.9 | n.d. | CPI | n.d. |

**Supplementary Table 4 | Detection of enzymes involved in sterol biosynthesis in the transcriptome of *Olavius algarvensis*.**

| Enzyme_ID | Contig | Querry | ID (%) | length | evalue | bitscore | assembly |
|---|---|---|---|---|---|---|---|
| ERG7 | TRINITY_DN47299_c3_g1_i19 | sp\|Q8BLN5\|ERG7_MOUSE | 61.165 | 721 | 0 | 894 | Oalg5ASA |
| ERG6 | TRINITY_DN46293_c3_g7_i1 | sp\|Q9LM02\|SMT1_ARATH | 49.158 | 297 | 8.50E-100 | 297 | OalgA5SA |
| ERG11 | TRINITY_DN47328_c5_g2_i2 | sp\|Q1JPY5\|CYP51_DANRE | 69.892 | 465 | 0 | 672 | OalgA5SA |
| ERG1 | TRINITY_DN49356_c3_g8_i1 | sp\|Q14534\|ERG1_HUMAN | 59.346 | 428 | 0 | 530 | OalgA5SA |
| ERG26 | TRINITY_DN40078_c3_g2_i1 | sp\|Q3ZBE9\|NSDHL_BOVIN | 60.117 | 341 | 1.02E-155 | 434 | OalgB8SA |
| ERG27 | TRINITY_DN48078_c2_g1_i1 | sp\|Q62904\|DHB7_RAT | 50 | 268 | 2.16E-83 | 251 | OalgA5SA |
| ERG24 | TRINITY_DN29949_c0_g1_i1 | sp\|Q14739\|LBR_HUMAN | 49.745 | 392 | 6.70E-137 | 397 | OalgB8SA |
| ERG25 | TRINITY_DN35202_c6_g4_i2 | sp\|Q5ZLL6\|MSMO1_CHICK | 64.789 | 284 | 6.74E-133 | 375 | OalgA8SA |
| partial_ERG3 | TRINITY_DN35917_c0_g1_i2 | sp\|O75845\|SC5D_HUMAN | 63.592 | 206 | 6.76E-101 | 301 | OalgB5SA |
| ERG2 | TRINITY_DN18361_c0_g1_i2 | sp\|P70245\|EBP_MOUSE | 45.249 | 221 | 3.50E-72 | 216 | verC3 |
| DHCR7a | TRINITY_DN28908_c0_g1_i1 | sp\|Q9LDU6\|ST7R_ARATH | 48.921 | 417 | 7.72E-137 | 394 | 4731 |
| DHCR24_4731_Verc3_consensus | | | | | | | |
| partial_DHCR24 | TRINITY_DN23155_c0_g1_i1 | sp\|Q5BQE6\|DHC24_RAT | 63.725 | 408 | 0 | 574 | 4731 |
| partial_DHCR24 | TRINITY_DN35798_c0_g2_i2 | sp\|Q5BQE6\|DHC24_RAT | 63.006 | 346 | 5.28E-175 | 488 | verC3 |

**Supplementary Table 5 | Detection of enzymes involved in sterol biosynthesis in the proteome of *Olavius algarvensis*.** FDR = False discovery rate, #PSMs = number of peptide spectral matches, #PUPs = number of protein unique peptides.

| Protein acession | Description | Found in proteome (filtered for 5% FDR) | Found in # of samples (out of 25) | q-value | #PSMs | #of PUP |
|---|---|---|---|---|---|---|
| ERG1_Host_330784_c8_seq1_40 | squalene monooxygenase homologue | No | - | - | - | - |
| ERG2_Host_282622_c0_seq2_5 | sterol C-8 isomerase homologue | Yes | 4 | 0.008 | 4 | 0 |
| ERG6_Host_316125_c3_seq1_6 | sterol methyltransferase homologue | Yes | 25 | 0 | 70 | 4 |
| ERG7_Host_333294_c4_seq2_44 | lanosterol synthase homologue | Yes | 3 | 0 | 4 | 2 |
| ERG11a_Host_334930_c0_seq4_30 | sterol C-14 demethylase homologue | Yes | 25 | 0 | 50 | 1 |
| ERG11b_Host_334930_c0_seq1_24 | sterol C-14 demethylase homologue | Yes | 12 | 0 | 16 | 0 |
| ERG24_Host_331074_c0_seq1_34 | sterol C-14 reductase homologue | Yes | 25 | 0 | 123 | 5 |
| ERG25_Host_335885_c4_seq2_8 | methylsterol monooxygenase homologue | No | - | - | - | - |
| ERG26a_Host_330893_c3_seq7_9 | Sterol-4-alpha-carboxylate 3-dehydrogenase | Yes | 17 | 0 | 25 | 1 |
| ERG27_Host_329156_c0_seq5_38 | 3-keto reductase homologue | No | - | - | - | - |
| ERG3_Host_326988_c0_seq1_35 | sterol C-5 desaturase homologue | No | - | - | - | - |
| DHCR24_4731_Verc3_concensus | sterol C-24 reductase homologue | No | - | - | - | - |
| DHCR7a_4731_TRINITY_DN28908_c0_g1_i1 | sterol C-7 reductase homologue | No | - | - | - | - |

**Supplementary Table 6 | The sterol profiles of all investigated *Olavius* and *Inanidrilus* species was dominated by sitosterol.**

| Species | Sampling location | Main sterol | other sterols | $C_{24}$-SMT |
|---|---|---|---|---|
| *Olavius algarvensis* | Sant'Andrea (Elba, Italy) | sitosterol | cholestrol | yes |
| *Olavius ilvae* | Sant'Andrea (Elba, Italy) | sitosterol | cholestrol | yes |
| *Olavius algarvensis* | Magaluf (Mallorca, Spain) | sitosterol | cholestrol | yes |
| *Olavius longissimus* | Carrie bow Cay (Belize) | sitosterol | cholestrol | yes |
| *Olavius tantulus* | Twin Cayes (Belize) | sitosterol | cholestrol | yes |
| *Inanidrilus mojicae* | Twin Cayes (Belize) | sitosterol | cholestrol | unknown |
| *Inanidrilus leukodermatus* | Carrie bow Cay (Belize) | sitosterol | cholestrol | yes |
| *Olavius sp.* | Okinawa (Japan) | sitosterol | cholestrol | yes |

**Supplementary Table 7 | $C_{24}$-SMT homologues were identified in the transcriptomes of all *Olavius* and *Inandrilus* species analyzed in this study**

| Species | Collection site | $C_{24}$-SMT | Target_ID | Type |
|---|---|---|---|---|
| *Inanidrilus leukodermatus* | Harrington Sound (Bermuda) | yes | TRINITY_DN9622 | transcript |
| *Inanidrilus sp.* FANTCC3 | Curlew Cay (Belize) | yes | TRINITY_DN8118 | transcript |
| *Inanidrilus sp.* NYSP | Carrie Bow Caye (Belize) | yes | TRINITY_DN8686 | transcript |
| *Inanidrilus sp.* ULE | Curlew Cay (Belize) | yes | TRINITY_DN41110 | transcript |
| *Olavius clavatus* | Lizard Island (Australia) | yes | g641.t1 | gene |
| *Olavius finitimus* | Twin Cayes (Belize) | yes | TRINITY_DN10503 | transcript |
| *Olavius ilvae* | Sant'Andrea (Elba, Italy) | yes | TRINITY_DN18930 | transcript and gene |
| *Olavius imperfectus* | Twin Cayes (Belize) | yes | TRINITY_DN8834 | transcript |
| *Olavius tantalus* | Twin Cayes (Belize) | yes | TRINITY_DN12053 | transcript |

**Supplementary Table 8 | $C_{24}$-SMT homologues are widely spread in annelids.** They were found in the transcriptomes of 9 *Olavius* and *Inanidrilus* species, in three deep-sea gutless tubeworm species and 17 gut-bearing annelid species from marine, limnic and terrestrial environments belonging to six different clades. Methylated sterols ($C_{28}$ and $C_{29}$) often account for an important part of the sterol profile of annelids. References: 1 **(Voogt, 1973a)**, 2 **(Voogt, 1973b)**, 3 (Marsh et al., 1990) , 4 **(McLaughlin, 1971)**, 5 (Petersen & Holmstrup, 2000), 6 (Hasan et al., 2012), 7 (Zipser et al., 1998), 8 (Naya & Kotake, 1967), 9 (Cerbulis & Wight Taylor, 1969), 10 (Albro et al., 1993), 11 (Mita et al., 2006), 12 (Guan et al., 2021), 13 (Kobayashi et al., 1973), 14 (Phleger et al., 2005), 15 (Rieley et al., 1996).
→ See attached excel file

**Supplementary Table 9 | $C_{24}$-SMT homologues are also present in sponges, rotifers and likely in mollusks.**
→ See attached excel file

**Supplementary Table 10 | Solvent gradient for high-resolution LC-MS/MS with a C30 column used to identify sterols.**

| %B | Time (min) |
|---|---|
| **0** | -2 (pre-run equilibration) |
| **0** | 2 |
| **16** | 5.5 |
| **45** | 9 |
| **52** | 12 |
| **58** | 14 |
| **66** | 16 |
| **70** | 18 |
| **75** | 22 |
| **97** | 25 |
| **97** | 32.5 |
| **15** | 33 |
| **0** | 34.4 |
| **0** | 36 |

Buffer A (60/40 ACN/$H_2O$, 10 mM ammonium formate, 0.1% FA) and Buffer B (90/10 IPA/ACN, 10 mM ammonium formate, 0.1% FA) were used at a flow rate of 350 µl $min^{-1}$

**Supplementary Table 11 | MS settings of the Q Exactive Plus Orbitrap (Thermo Fisher Scientific) equipped with a HESI probe and a Vanquish Horizon UHPLC System (Thermo Fisher Scientific).**

| $MS^1$ | C30 settings | C18 settings |
|---|---|---|
| Resolution | 70,000 | 70,000 |
| AGC target | 5e5 | 3e6 |
| Max IT [ms] | 65 | 200 |
| Scan range [*m/z*] | 150–1500 | 100–1000 |
| **$MS^2$** | **DDA** | **DIA** |
| Resolution | 35,000 | 35,000 |
| AGC target | 1e6 | 2e5 |
| Max IT [ms] | 75 | auto |
| Loop count | 8 | 1 |
| Dynamic exclusion [sec.] | 30 | NA |
| Isolation windows (pos.) [Da] | 1 | 0.4 |
| Isolation windows (neg.) [Da] | 1 | NA |
| NCE | 30 | 30 |

**Supplementary Table 12 | List of the enzymes isolated from model organisms used as query to assess the ability of gutless annelids to synthesize sterols.**

| Enzyme_name | animal | plant | fungi |
|---|---|---|---|
| **SQE/ERG1** | Q14534 | Q9SM02 | Q92206 |
| **LAS/CAS/ERG7** | P48449 | P38605 | P38604 |
| **CYP51/ERG11** | Q16850 | Q9SAA9 | P10614 |
| **LBR/FK/ERG24** | O76062<br>Q14739 | P32462 | Q9LDR4 |
| **MSMO1/SMO/ERG25** | Q15800 | Q8L7W5<br>Q1EC69<br>Q9ZW22<br>Q8VWZ8<br>F4JLZ6 | O59933 |
| **NSDHL/3β-HSD/ERG26** | Q15738 | A9X4U2<br>Q67ZE1<br>Q9FX01 | P53199 |
| **HSD17B7/?/ERG27** | P56937 | - | Q12452 |
| **EBP/HYD1/ERG2** | Q15125 | O48962 | P32352 |
| **SC5DL/DWF7/ERG3** | O75845 | Q39208<br>Q9M883 | P32353 |
| **DHCR7/DWF5/-** | Q9UBM7 | Q9LDU6 | - |
| **DHCR24/DIM/ERG4** | Q15392 | Q39085 | P25340 |
| **-/CYP710A/ERG5** | - | O64697<br>O64698<br>Q9ZV28<br>Q9ZV29 | P54781 |
| **-/SMT/ERG6** | - | Q9LM02<br>Q39227<br>Q94JS4 | P25087 |
| **-/CPI/-** | - | Q9M643 | - |

| Enzyme_name | animal | plant | fungi |
|---|---|---|---|
| **SQE/ERG1** | Q14534 | Q9SM02 | Q92206 |
| **LAS/CAS/ERG7** | P48449 | P38605 | P38604 |
| **CYP51/ERG11** | Q16850 | Q9SAA9 | P10614 |
| **LBR/FK/ERG24** | O76062<br>Q14739 | P32462 | Q9LDR4 |
| **MSMO1/SMO/ERG25** | Q15800 | Q8L7W5<br>Q1EC69<br>Q9ZW22<br>Q8VWZ8<br>F4JLZ6 | O59933 |
| **NSDHL/3β-HSD/ERG26** | Q15738 | A9X4U2<br>Q67ZE1<br>Q9FX01 | P53199 |

| | | | |
|---|---|---|---|
| **HSD17B7/?/ERG27** | P56937 | - | Q12452 |
| **EBP/HYD1/ERG2** | Q15125 | O48962 | P32352 |
| **SC5DL/DWF7/ERG3** | O75845 | Q39208<br>Q9M883 | P32353 |
| **DHCR7/DWF5/-** | Q9UBM7 | Q9LDU6 | - |
| **DHCR24/DIM/ERG4** | Q15392 | Q39085 | P25340 |
| **-/CYP710A/ERG5** | - | O64697<br>O64698<br>Q9ZV28<br>Q9ZV29 | P54781 |
| **-/SMT/ERG6** | - | Q9LM02<br>Q39227<br>Q94JS4 | P25087 |
| **-/CPI/-** | - | Q9M643 | - |

**Supplementary Table 13 | Details of the sequences, plasmid, and *E. coli* cells used for the heterologous gene expression experiments.**

| Gene Name | Sequence | Length | Restriction Sites to Keep | Cloning Vector | Comment | Expression host | ID |
|---|---|---|---|---|---|---|---|
| AraTh_Q9LM02 | MDLASNLGGKIDKSDVLTAVEKYEQYHVF<br>HGGNEEERKANYTDMVNKYYDLATSFYE<br>YGWGESFHFAQRWKGESLRESIKRHEHF<br>LALQLGIPGQKVLDVGCGIGGPLREIARFS<br>NSVVTGLNNNEYQITRGKELNRLAGVDKT<br>CNFVKADFMMPFPENSFDAVYAIEATCHA<br>PDAYGCYKEIYRVLKPGQCFAAYEWCMT<br>DAFDPDNAEHQKIGEIEIGDGLPDIRLTTKC<br>LEALKQAGFEVIWEKDLAKDSPVPWYLPL<br>DKNHFSLSSFRLTAVGRFTKNMVKILEYIR<br>LAPQGSQRVSNFLEQAAEGDGGRREIFT<br>PMYFFLARKPE | 1023 bp | NheI/XhoI | pET-28a(+) with restriction sites NheI/XhoI | His- Tag at the N-terminal side only | *E. coli* C41(DE3)pLysS | T-3784 |
| OclaLIZ1_g641.t1 | MTSVEKPSITEILRPLQAKSPSSVESTADG<br>YLRYFEREQQSKDDKVDEEDLDAEATDR<br>RRQDAVTVTNAYYDLATDFYEYGWGDFH<br>FAVLKPEESREHSFAKHEYFLAMKLGLKA<br>GDTVLDIGCGIGGPARHIASLSEANVIGMN<br>INDYQLSRARILTEKAKLDHLCSFVKADYN<br>HMPYGDGHFDVYAIEATCHSPTLLSVYSE<br>VFRVLKPGGMFAVYEWIMTDKYNPTDPY<br>HKKLKADILEGDGLPDIVTAPQAVVAARQA<br>GFEVLESRDRALEPGLPWNVLQARWTLS<br>DIKITPLGRWATHVMLAVLETVHLAPRGAV<br>KVHRTLCKGADALAAAGVEGIFSPMYLLV<br>LRKPRD | 1089 bp | NheI/XhoI | pET-28a(+) with restriction sites NheI/XhoI | His- Tag at the N-terminal side only | *E. coli* Lemo21(DE3) | T-3787 |
| OalgB8SA | MNSIEKPSITGILRPLHGKAAVESTADGYL<br>RYFDSDSPPRDDAADADDSDAEAVERRR<br>KNALAVTNAYYDLATDFYEYGWGEAFHFA<br>VLKPEESREHSFAKHEYFLGMKLGLKAGD<br>TVLDIGCGIGGPARHIASLSEANVIGLNIND<br>YQLSRARILTEKAKLDHLCSFVKADYNHMP<br>YGEGHFDKVYAIEATCHSPSLCSVYSEVF<br>RVLKPGGLFALYEWIMTDTYNPTDPYHKK<br>LKADILEGDGLPDLASVPQVLTAARQAGF<br>EVVESRDRALEPGVPWYTVLQARWTLSDI<br>KITPFGRWATHLMLAVLETVRLAPRGSVK<br>VHRTLCKGADALAAAGAEGIFSPMYLLVL<br>RKPSK | | NheI/XhoI | pET-28a(+) with restriction sites NheI/XhoI | His- Tag at the N-terminal side only | *E. coli* Lemo21(DE3) | T-3909 |

**Supplementary Table 14 | List of the different sterol substrates tested with the animal $C_{24}$-SMTs.** *Olavius algarvensis* $C_{24}$-SMT (Oalg_SMT), *Olavius clavatus* $C_{24}$-SMT (Oclav_SMT) and *Arabidopsis thaliana* $C_{24}$-SMT (Atha_SMT1). Green shading indicates that the substrate can be used by the enzyme tested. Orange shading indicates that no methylated product was detected at the end of the incubation between the set of enzyme and the substrate tested.

| Substrate | Enzyme | | | Rationale for testing the substrate |
|---|---|---|---|---|
| | Oalg_SMT | Oclav_SMT | Atha_SMT1 | |
| Lathosterol | | | | Cholesterol intermediate |
| 7-dehydrocholsterol | | | | Cholesterol intermediate |
| Desmosterol | $1^{st}$ methylation | $1^{st}$ methylation | | Cholesterol intermediate |
| Cholesterol | | | | Cholesterol intermediate |
| Zymosterol | $1^{st}$ methylation | $1^{st}$ methylation | | Substrate of fungal $C_{24}$-SMT and cholesterol intermadiate |
| Eburicol | | | | *Pneumocystis carninii*, substrate for $2^{nd}$ methylation |
| Lanosterol | | | | Substrate of fungal $C_{24}$-SMT |
| Cycloartenol | | | $1^{st}$ methylation | Substrate of the first methylation in plants |
| Campesterol | | | | Potential intermediate of *O. algarvensis* sitosterol biosynthesis |
| 24-methylene-cholesterol | $2^{nd}$ methylation | $2^{nd}$ methylation | | Potential intermediate of *O. algarvensis* sitosterol biosynthesis |

**Supplementary Table 15 | Solvent gradient for high-resolution LC-MS/MS with C18 column.**

| %B | Time (min) |
|---|---|
| **50** | -10 (pre-run equilibration) |
| **50** | 3 |
| **87.5** | 9 |
| **90** | 15 |
| **100** | 21 |
| **100** | 30 |
| **50** | 32 |

*Solvent A (MiliQ, 0.1% FA) and Solvent B (90:10 ACN:MiliQ, 0.1% FA) were used at a flow rate of 200 µl min$^{-1}$.*
